# NSCLC patients with oligo-metastatic brain disease show an altered CD4 T-cells immune profile

**DOI:** 10.1101/2024.04.10.588829

**Authors:** Mais Alsousli, Cecile L. Maire, Andras Piffko, Jakob Matschke, Laura Glau, Merle Reetz, Svenja Schneegans, Gresa Emurlai, Benedikt Asey, Alessandra Rünger, Sven Peine, Jolanthe Kropidlowski, Jens Gempt, Markus Glatzel, Manfred Westphal, Eva Tolosa, Katrin Lamszus, Klaus Pantel, Simon A. Joosse, Malte Mohme, Harriet Wikman

## Abstract

**Background:** Lung cancer is the leading cause of cancer-related deaths worldwide, with brain metastasis (BM) occurring in 40% of advanced non-small cell lung cancer (NSCLC) patients. In 15% of these patients, the brain is the only affected organ (oligo-metastasis), corresponding to improved prognosis compared to widespread disease. Thus far, it is unknown if the metastatic dissemination to the brain without systemic metastases is a consequence of the immune system’s ability to control systemic tumor outgrowth.

**Methods:** Here, we investigated the local and peripheral immune cell composition in NSCLC BM patients, and identified new immune patterns related to the occurrence of brain metastases either as oligo- or poly-metastatic disease.

**Results:** The multi-parametric immune phenotyping of peripheral blood revealed a downregulation of KLRG1 in CD8^+^ T-cells and an increase in CD4^+^ T_H_17 cells and elevated IL-17 levels in the blood of all NSCLC BM patients compared to healthy individuals. In addition, BM patients CD4^+^ T cells showed less CD73 expression with reduced effector memory differentiation. Furthermore, we observed less intra-tumoral infiltration in tumor tissues and a distinctive CD4+ T-cell profile in oligo-synchronous BM, both in the tumor microenvironment and peripheral blood compared to poly-metastatic BM patients. Moreover, 5′-ectonucleotidase CD73 was significantly upregulated in CD4 and T-regulatory cells of oligo-synchronous BM.

**Conclusions:** These results indicate that oligo-synchronous BM exhibits a more pronounced alteration in the CD4 T-cell immune profile both locally at the tumor site and systemically.

**Key Points:**

- BM patients exhibit a skewed systemic immune profile, characterized by downregulation of KLRG1 in CD8^+^ and induction of T_H_17/IL-17 axis and CD73 in CD4^+^ T-cells.
- Oligo-synchronous BM displayed a distinct CD4^+^ T-cell profile in both TME and peripheral blood.

**Importance of the Study:** This study presents a novel insight into immune profiles of brain metastasis types in NSCLC patients. Examining tissues and PBMCs sheds light on the disease and uncovers unique immune responses within distinct brain metastasis patterns. This research offers valuable knowledge for improved understanding and identifying potential prognosis markers.

## Introduction

Lung cancer is the leading cause of cancer-related deaths worldwide, killing more than colon, breast, and prostate cancers combined ^1^. This high rate of mortality is attributed to the early and rapid metastasis to multiple organs, with the brain being affected in approximately 40% of advanced non-small cell lung cancer (NSCLC) patients ^2^. Therefore, brain metastases (BM) have become a major limitation of life expectancy and quality of life for many individuals. Consequently, developing effective management strategies for brain metastases is highly important.

The metastatic spread of primary tumors is neither random nor merely affected by anatomical location. However, the exact mechanisms responsible for the patterns of metastasis remain unclear. Among others, host-related factors such as immunological properties and target-organ microenvironment are suspected to play an important role ^3^. A distinct feature of NSCLC is the frequent occurrence of oligo-metastatic disease in the brain. In fact, more than 15% of all metastatic lung adenocarcinomas exclusively affect the brain as the only organ involved, either at the time of primary diagnosis (oligo-synchronous) or after primary diagnosis (oligo-metachronous) ^2^. Clinical data suggest that oligo-brain metastatic disease, often accompanied by a favorable prognosis, represents a distinct form of cancer spread compared to a disease that spreads uncontrolled to multiple sites (poly-metastatic) ^4–6^. Several prospective studies have demonstrated significantly improved progression-free or overall survival in oligo-metastatic NSCLC patients who received stereotactic body radiotherapy, as compared to maintenance therapy alone ^7–9^.

A deregulated immune system is widely recognized as one of the hallmarks of cancer, in which both positive and negative interactions between immune and neoplastic cells play a major role in malignant progression ^10^. Due to the unique properties and immune contexture of the brain microenvironment ^11,12^, we hypothesized that the phenotype and activation pattern of immune cells is different in the tumor microenvironment as well as in the peripheral blood when metastases home to the brain. In the study presented here, we aimed to discern immune cell populations in both brain metastatic tissue and peripheral blood to map the immunological landscape in NSCLC patients affected with BM, as well as to identify new immune patterns related to the occurrence of brain metastases either as oligo- or poly-metastatic disease. Beyond understanding the biological basis for different metastatic patterns in BM, such approaches can provide new insights for immunomodulatory treatment strategies targeting specific metastasis induced inflammatory states.

## Materials and Methods

### Patient cohorts

All patients included in this study were operated for lung cancer brain metastases at the Department of Neurosurgery at University Hospital Hamburg-Eppendorf (UKE) between 2014 and 2020. Patients with known driver mutations in *EGFR*, *ALK*, or *ROS* were excluded before recruitment. The clinical characteristics of 69 NSCLC patients are summarized in Table 1. Of these, 54 patients were included in the peripheral blood analyses and 56 in tissue phenotyping including 28 non-overlapping samples (Supplemental tables 1 and 2). This study was approved by the local ethics council of the Hamburg chamber for physicians under numbers Nr. PV5392 & Nr. PV4904 and was performed in accordance with the Helsinki Declaration of 1975.

**Table 1:** Patient characteristics.

| Characteristics |  | All BM | Oligo-synchronous | Oligo-metachronous | Poly-metastatic | P-value |
| --- | --- | --- | --- | --- | --- | --- |
| **Samples, n (%)** |  | 69 | 23 (33.3) | 20 (29) | 26 (37.7) | 0.368 |
| **Gender, n (%)** | F | 38 (55.1) | 10 (43.5) | 11 (55) | 17 (65.4) | 0.331 |
|  | M | 31 (44.9) | 13 (56.5) | 9 (45) | 9 (34.6) |  |
| **Age at PD (y)** | Range | 33 – 78 | 45 – 78 | 34 – 76 | 33 – 77 | 0.48 |
|  | Mean | 59.3 | 61.2 | 59.2 | 57.7 |  |
| **Age at BM-OP (y)** | Range | 34 – 83 | 45 – 83 | 34 – 78 | 35 – 78 | 0.422 |
|  | Mean | 60.2 | 61.6 | 61.3 | 58.1 |  |
| **Histology, n (%)** | AC | 63 | 21 | 16 | 26 | 0.141 |
|  | SQ | 3 | 1 | 2 | 0 |  |
|  | LCNEC | 3 | 1 | 2 | 0 |  |
| **Time btw PD & BM OP (months)** | Range | 0 – 69 | 0 | 2 – 69 | 0 – 30 | 0.965 |
|  | Mean | 10.2 | 0 | 24.3 | 5.4 |  |
| **Neoadjuvant treatment** | none | 41 | 22 | 1 | 17 | <0.0001 |
|  | IT | 4 | 0 | 1 | 3 |  |
|  | RCT | 24 | 1 | 18 | 6 |  |
| **Survival (m)** | alive | 20 | 6 | 4 | 10 | 0.380 |
|  | dead | 49 | 17 | 16 | 16 |  |
| **FUP after BM OP (m)** | Range | 0 - 83 | 2 - 65 | 0 - 59 | 0 - 83 | 0.678 |
|  | Mean | 17.7 | 20.2 | 17.4 | 15.2 |  |

BM: brain metastases, PD: Primary diagnosis, BM-OP: brain metastases operation, F: female. M: male, AC: adenocarcinoma, SQ: Squamous cell carcinoma, LCNEC: Large cell neuroendocrine carcinoma, TN: treatment-naïve, IT: immunotherapy, RCT: Radio-/Chemotherapy

### Immunohistochemical assays

Immunohistochemical staining (IHC) was performed on 2µm thick sections of formalin-fixed, paraffin-embedded (FFPE) tissue samples of BM (n=56) from the Department of Neuropathology, UKE). Antibodies used for IHC were CD3 (clone SP7; 1:100; M3074, Zytomed), CD8 (clone SP16; 1:500; C1008C01, DCS), CD4 (clone 4B12; 1:50; M731001-2, DAKO), FOXP3 (clone D2W8E; 1:100; 98377, Cell Signaling), CD68 (clone KP1; 1:100; M087629-2, DAKO), and Ki67 (clone SP6; 1:750; 275R-15, Cell Marques). All staining was performed on a Ventana benchmark XT autostainer following the manufacturer’s recommendations. Percentages of positively stained cells were evaluated in two regions, intra-tumoral (within the solid tumor tissue area) and peri-tumoral (the adjacent area around the tumor tissue). Immunohistochemical evaluation was performed semi-quantitatively by a pathologist as described earlier ^13^, briefly, for CD3, CD8, CD4, CD68, and Ki67, a score of 0 (negative: no stained cells), 1 (low: <10%), 2 (moderate:10-40%) or 3 (high >40%) was given based on the estimation of positive stained immune cells out of total cells in the investigated intra-tumoral or peri-tumoral region (n=56). For the statistical analysis moderate and high patients were combined (as high) and negative and low (as low). Since the number of FoxP3^+^ cells was relatively low in all BM groups, samples were scored as negative (0%) or positive (≥1%) in both intra-tumoral and peritumoral regions (n=60).

### Immune cell isolation

Peripheral blood was collected into 7.5ml EDTA-containing tubes before surgical removal of the tumor. In addition to 54 patients, control samples were obtained from 20 anonymized, age-matched healthy donors (HD) from the Department of Transfusion Medicine, University Hospital Hamburg-Eppendorf (Hamburg, Germany). PBMC were isolated within 2 hours of blood draw using Ficoll^®^ gradient centrifugation as described before ^14^ and stored in RPMI/10% DMSO (P04-17500) at −80°C until further usage.

### Multicolor flow cytometry

Frozen PBMCs were thawed in a water bath at 37°C, washed with 4°C cold 10% FBS in RPMI, resuspended in a cold medium (RPMI, 10% FCS and DNAse (1:1000) (4°C), and counted using Vi-CELL^®^ XR Cell Viability Analyzer (383556, Beckman Coulter). Five different panels (43 antibodies) were used to analyze T-cell exhaustion, T-cell differentiation, T helper cell subsets, T-cell metabolism, and cytokines secretion (Supplemental table 3).

For cell surface staining, samples were resuspended in flow-cytometry staining buffer (eBioscience) with Fc-block (Miltenyi Biotec), then stained with the antibody cocktails at room temperature for 45 minutes in the dark. For intracellular staining (cytokines), after 4 hours of incubation at 37°C and 5% CO_2_, a stimulation mix (Phorbol 12-Myristate 13-Acetate (PMA) (1µg/ml; P1585, Sigma), Ionomycin calcium salt (1mg/ml; 10634, Sigma), Brefeldin A Solution 1000X (3mg/ml; 00-4506, Invitrogen) and X-Vivo 15 serum-free (881024, Biozym) was added to each sample, and cells were incubated again for another 5 hours at 37°C and 5% CO_2_. Samples were then washed, resuspended in flow-cytometry staining buffer, and stained with a surface antibody cocktail for 10 minutes in the dark. After further washing steps, cells were fixed and permeabilized before staining with the cytokine (intracellular) antibody cocktail for 30 minutes in the dark at room temperature. Following the incubation, cells were washed and resuspended in a flow-cytometry staining buffer. Analysis was performed on a BD LSR Fortessa flow cytometer. Data were exported as .fcs files and manual gating was carried out using the FACSDiva software (version 9.1 Becton Dickinson). The gating strategies are shown in Supplementary figures 1-4. Samples with data from less than 1000 living T cells were excluded from the analyses.

### Statistical analysis

Data analysis was performed with In-Silico Online v2.3.1, R version 4.1.3 (http://in-silico.online), and GraphPad Prism Software (v9.5.0, GraphPad Inc., Boston, MA, USA). Associations between independent nominal data were calculated using Fisher’s exact test, whereas dependent data were assessed using McNemar’s test. Kruskal-Wallis and Wilcoxon’s tests were performed to calculate the significance of the differences between medians, and the significance of the mean difference was calculated using ANOVA. An alpha level of 0.05 was used to determine statistical significance and was corrected for multiple testing where appropriate. Survival analyses were performed using the Kaplan-Meier curves and log-rank test. 48 patients were included in the survival analyses as those patients who had no follow-up or were deceased during the perioperative period of 3 Months were excluded. Overall survival (OS) was estimated from the date of brain surgery until death or censored at the last follow-up. Hazard ratios (HR) were estimated using the Cox proportional regression model.

For panel 2 of flow-cytometry, we performed UMAP unsupervised clustering analysis, as compensated and normalized flow data were used to generate UMAP dimensionality reduction (using the R-package “umap”) and PhenoGraph clustering of all groups together (using R-package “Rphenograph”). The heatmap of marker expression was calculated as the median fluorescence intensity of each marker for each phenoGraph cluster. Comparisons were calculated with the Kruskal-Wallis test.

## Results

### Patients’ characteristics

Sixty-nine NSCLC patients with BM were recruited for this study (Table 1, Supplemental Tables 1 and 2). The patients were divided into two main groups: 1) NSCLC patients with BM only (oligo-metastatic; lymph node dissemination allowed) and 2) NSCLC patients with metastases in the brain and other extra-cranial organs (poly-metastatic). The oligo-metastatic group was further subdivided into: 1a) patients with brain metastasis at the time of primary diagnosis (synchronous–oligo metastasis) and 1b) patients with brain metastases after primary diagnosis (metachronous–oligo metastasis). Regarding the main clinical determinants of the three groups, neoadjuvant treatment was the only significant variable difference between the groups (P<0.0001, Table 1). Namely, most patients with oligo-metachronous metastases received pre-treatment (19/20), whereas only one patient with oligo-synchronous disease (1/23) and nine patients (9/26) with poly-metastatic disease only received neoadjuvant chemotherapy.

### Immune profiling of NSCLC brain metastasis

To establish a brain metastasis-associated immune profile, the spatial distribution of infiltrating immune cells was evaluated in peri-tumoral and intra-tumoral regions of BM tissues from NSCLC patients by staining CD3, CD8, CD4, FoxP3, and CD68 (Figure 1a and supplementary figure 5). In addition, tumor cell proliferation was determined by Ki67 (supplementary figure 5).

**Figure 1:**
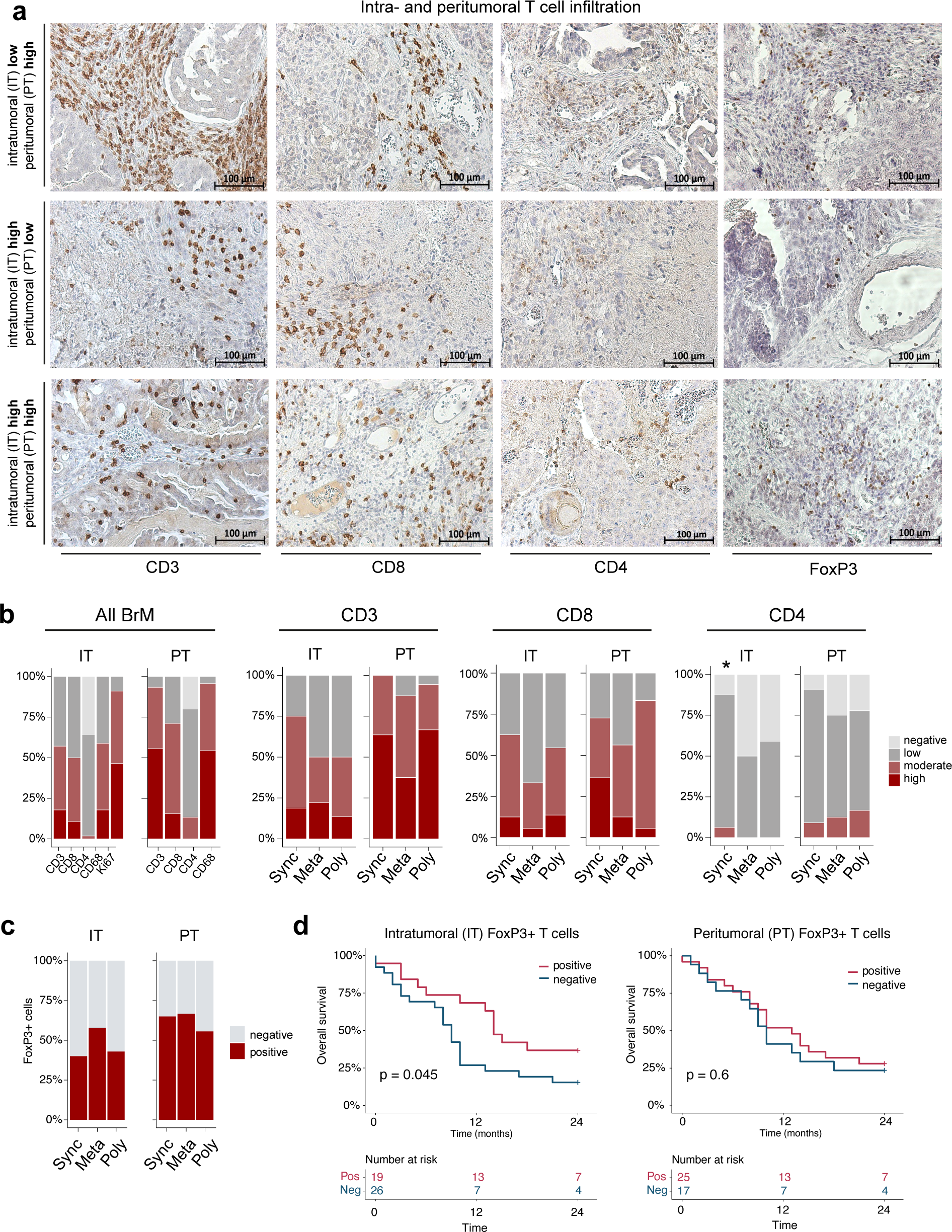
Immunohistochemical detection and estimation of positive staining for CD3, CD8, CD4, and FoxP3 on TILs in brain metastatic tissues. Representative stainings are shown for both intra-tumoral (IT) and peri-tumoral (PT) regions. (a.) Representative images of CD3, CD8, CD4, and FoxP3 staining results respectively in IT and PT regions (high and low infiltration). Magnification: x20. (b.) Comparison of the staining scores for CD3, CD8, and CD4 in all groups combined, and between oligo-synchronous BM (Sync), oligo-metachronous BM (Meta), and poly BM (Poly) individually in both IT and PT regions. (c.) Comparison of the staining scores for FoxP3 between BM groups in IT and PT regions. (d.) Kaplan-Meier survival curves (24m) based on FoxP3 expression in IT and PT regions. BM: brain metastasis, Sync: oligo-synchronous BM, Meta: oligo-metachronous BM, and poly: poly metastasis, negative: no stained cells, low: <10%, moderate: 10-40%, and high: >40%

Overall, a significantly higher frequency of CD3^+^ T-cell infiltration and CD68^+^ microglia/macrophages could be detected in peri-tumoral regions (93.3 and 95.7% moderate/high inflitration, respectively) compared to the core of the BM (intra-tumoral) (57.1 and 58.9%) (P<0.0001 for both markers, Fisher’s exact test). Furthermore, within the CD3^+^ T-cell population, a significantly greater proportion of CD8^+^ cells moderate & high infiltration) was observed in both intra-tumoral and peri-tumoral lesions (50.0 and 71.1% respectively) compared to CD4^+^ cells (1.8 and 13.3% respectively) (P<0.0001 in both regions, McNemar’s test). The average proliferation rate of the tumors was 44% (range 5-90%). The proliferation rate did not correlate with the immune cell profiles (Figure 1b). Taken together, different types of immune cells infiltrate the brain in NSCLC BM, however, these cells are found more commonly in the peri-tumoral area compared to intra-tumoral regions.

When analyzing the oligo- and poly-metastatic cases separately, IHC staining did not reveal any significant differences in the number of CD3^+^ or CD8^+^ cells between the groups in either region (Figure 1b). However, patients with oligo-synchronous BM, demonstrated a significantly higher infiltration of CD4^+^ T-cells in the intra-tumoral region compared to poly- and oligo-metachronous BM patients (P=0.044, G-test). Specifically, within the latter two groups, 50% and 60% of samples had detectable CD4^+^ T-cells within the tumor tissues, whereas 87.5% of oligo-synchronous cases showed CD4^+^ T-cell infiltration (Figure 1b). No significant differences were observed in the number of microglia/macrophages (CD68^+^ cells), or in Ki67-positive tumor cells among the three BM groups (Supplementary Figure 5).

As the tumor-infiltrating CD4^+^ cell population was found to be elevated in the oligo-synchronous BM, we further investigated the frequency of regulatory T-cells (Tregs) defined by FoxP3 expression (Figure 1a). Comparisons among the BM groups revealed no statistically significant differences in Tregs, neither intra-tumoral nor peri-tumoral (P=0.524 and 0.831, respectively, Fisher’s exact test; Figure 1c). However, the frequency of FoxP3^+^ intra-tumoral infiltrating cells showed a significantly positive correlation with overall survival (P=0.045, log-rank test), but not in the peri-tumoral region (P= 0.600, log-rank test), for all patients (Figure 1d). The presence of intra-tumoral FoxP3 expressing cells was independently associated with survival in multivariable analysis (HR:0.415, 95% CI:0.210-0.821, P= 0.011; Cox proportional hazard ratio). No significant difference in survival was observed related to the number of infiltrated CD3, CD8, CD4, or CD68 cells in the intra- or peri-tumoral regions (p>0.05, log-rank test) (Supplemental figure 6).

### Systemic immunomodulation in patients with NSCLC compared to healthy donors

We performed flow-cytometry analyses on the mononuclear cell fraction of peripheral blood samples obtained from NSCLC BM patients and compared the T- and NK cell immunophenotypes to those of age-matched healthy individuals using five different multicolor antibody staining cocktails (Figure 2a). NK cells and T-cell profiles were investigated with focus on T-cell exhaustion, T-helper cell differentiation subsets, T-cell metabolism, and cytokine secretion, focusing on the expression of activation and exhaustion markers, such as PD-1, KLRG1, TIGIT, CD39, and CD73. The heatmaps depicts heterogeneous expression of these markers among the NSCLC patients compared to healthy donors (Figure 2b). For activation and exhaustion markers, the frequency of 5′-ectonucleotidase CD73, which converts adenosine monophosphate to the immuno-inhibitory adenosine, expressed by CD8^+^ and CD4^+^ T-cells was significantly upregulated in the cancer patients compared to healthy individuals (Figure 2c, P=0.022 and P=0.0001, respectively Welch’s t-test). No significant difference was observed for the frequency of cells expressing the classical immune checkpoint PD-1, although a trend in the CD8^+^ T cell compartment was measurable (Figure 2c, P=0.063, Welch’s t-test). The co-inhibitory receptor KLRG1 is typically expressed on late-differentiated CD8^+^ and NK cells, indicating immunosenescence ^15^. In the blood of our patient cohort, the mean percentage of KLRG1 expressing CD8^+^ cells was 32.7% and 52.9% in healthy individuals (P<0.0001, Welch’s t-test; Figure 2c). On the other hand, we observed lower levels of CD4^+^ early memory T-cells (CD45RA^-^, CD27^+^, CD28^+^, CCR7^-^) in cancer patients (P= 0.024, Welch’s t-test, Figure 2c). Furthermore, a significant mean difference in Treg expression TIGIT was observed between the cancer patients and healthy individuals (P=0.001, Welch’s t-test, Figure 2d), implying a less immunosuppressive functional state during NSCLC brain metastasis ^16^. No significant differences in the presence of NK cells expressing activation/exhaustion markers between the blood of cancer patients and healthy individuals could be observed. Taken together, this analysis shows a mixed phenotype, as potentially tumor-induced immunosuppressive molecules such as CD73 and PD-1 are upregulated on peripheral T-cells, while others, like TIGIT and KLRG1, are reduced in NSCLC BM patients.

**Figure 2:**
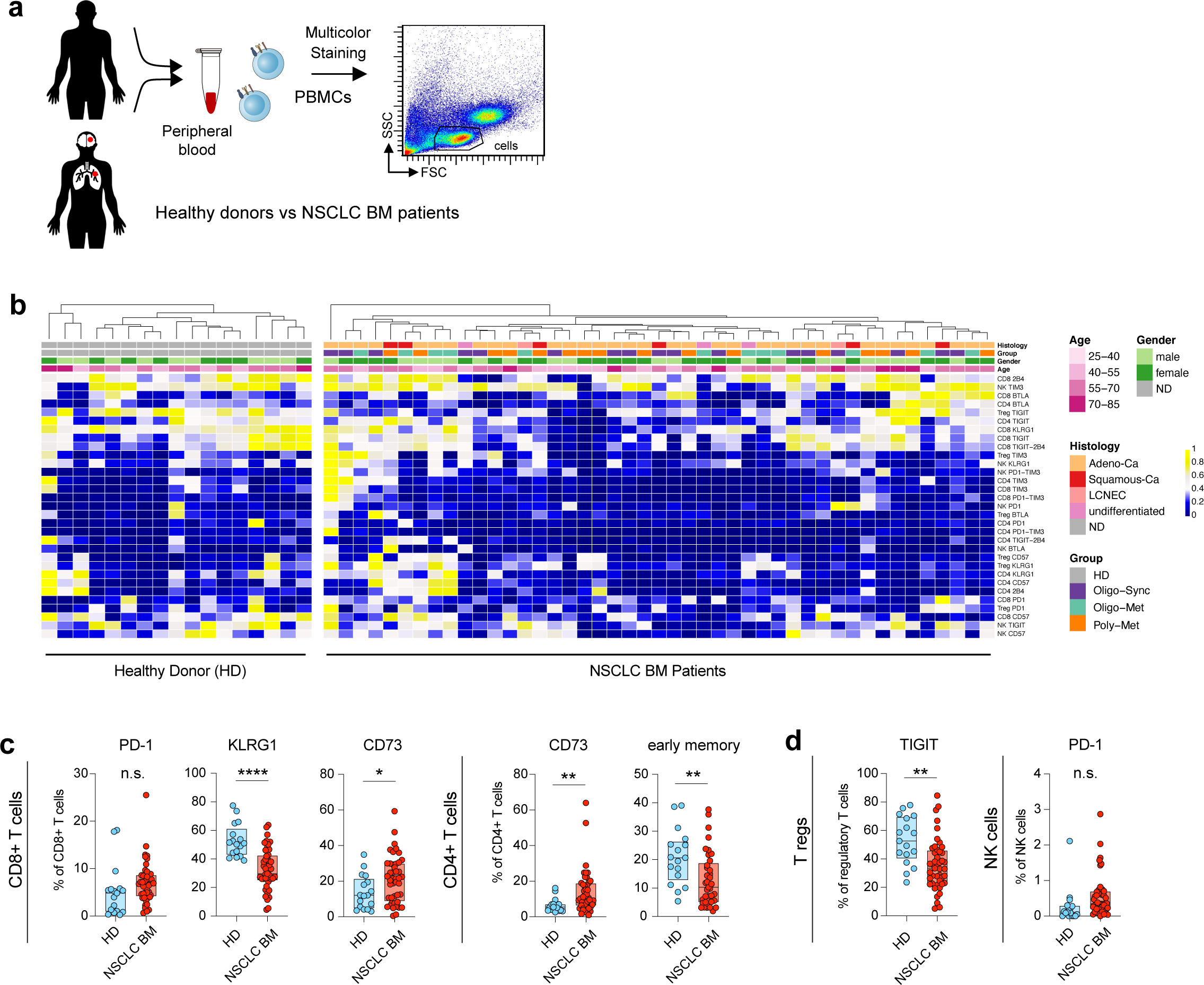
Flow cytometric immunophenotyping of T-cell activation and exhaustion marker expression. (a.) Flow cytometry analysis of peripheral blood from NSCLC BM patients and age-matched healthy donors (HD). (b.) Heatmap representing the median expression of activation/exhaustion markers in NSCLC BM and HD samples. (c. and d.) Box plots depicting expression of PD-1, KLRG1, CD73, early memory and TIGIT in different tumor-infiltrating T-cell populations. Mean ± SEM, t test. P values are defined as * < 0.05; ** < 0.01; and *** < 0.001.

Next, we analyzed the composition of the peripheral CD4^+^ T helper cell subsets in NSCLC BM patients compared to healthy individuals. While no differences between the study participants were observed in terms of the classical T_H_1 and T_H_2 subsets (Figure 3a and b), we found a significantly higher frequency of pro-inflammatory T_H_17 T-cells (CD4^+^, CD45RA^-^, CCR6^+^, CCR4^+^, CXCR3^-^, CD161^+^; Figure 3b; P=0.004, Welch’s t-test) among the cancer patients compared to healthy individuals. T_H_17 T-cells produce the pro-inflammatory cytokine IL-17 and their ultimate role in cancer is incompletely understood with both pro and antitumor functions described ^17,18^. However, this finding is in line with our observation of an increased activation of the T_H_17 axis, and thus a potentially pro-inflammatory peripheral immune environment in NSCLC BM patients.

**Figure 3:**
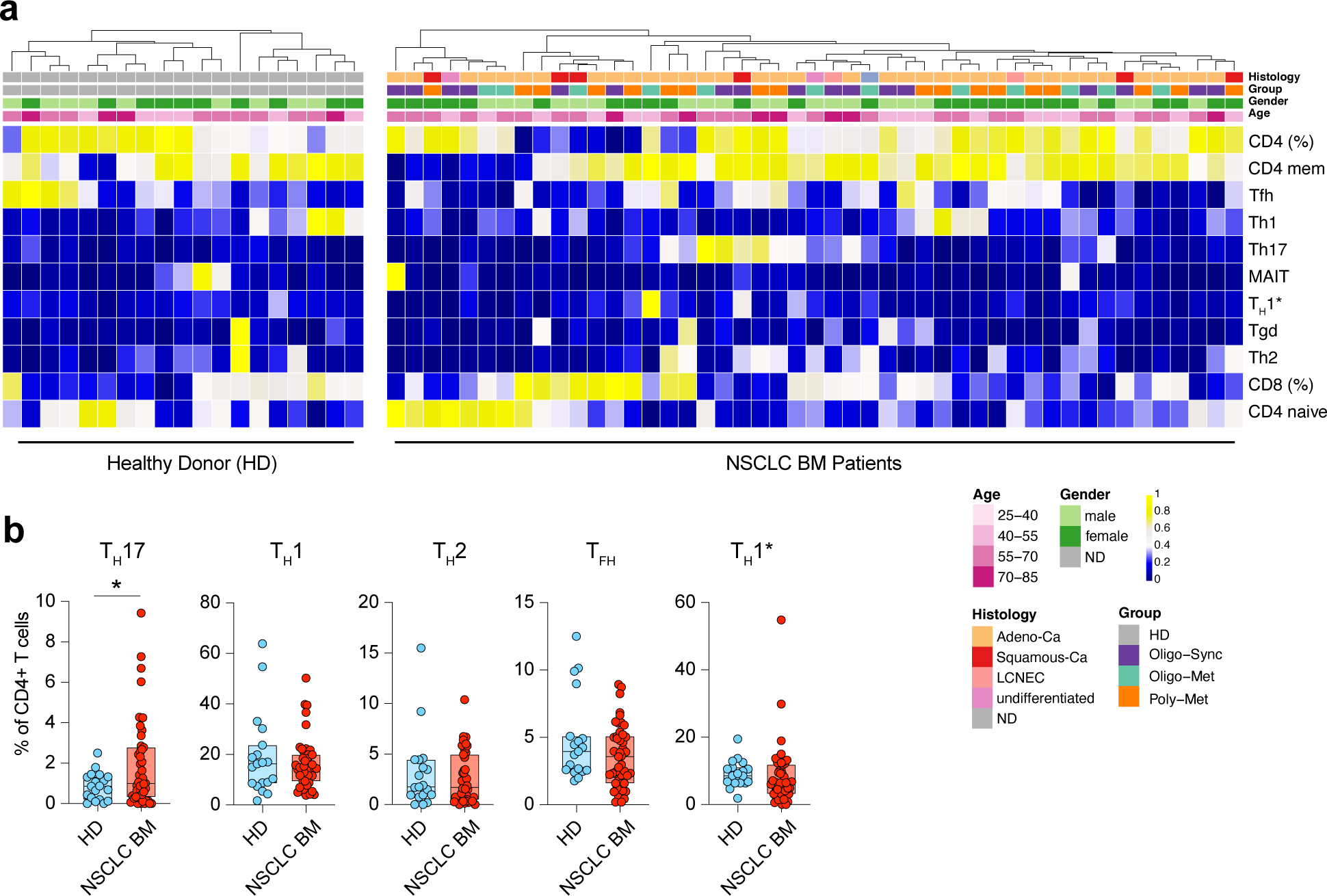
Flow cytometric immunophenotyping of Th subsets. (a.) Heatmap representing the median expression of Th subsets in NSCLC BM and healthy donors HD samples. (b.) Box plots depicting expression of TH1, TH2, TH17, TH1*, and TFH cells in CD4 T-cell populations. Mean ± SEM, t test. P values are defined as * < 0.05; ** < 0.01; and *** < 0.001.

To follow up on this finding, we conducted intracellular cytokine staining on T- and NK cells from the peripheral blood of NSCLC BM patients in comparison to healthy individuals. PBMCs were stimulated using PMA/ionomycin, and we assessed the accumulation of hallmark cytokines intracellularly using flow cytometry. Among the NSCLC patients, we found that 14/28 (50%) of NSCLC BM patients showed a reduced overall cytokine production (Figure 4a), while the other half of the patients could be divided into a group of predominantly TNFa and IFNy producers (9/14), and a second subgroup (5/14) of patients with a poly cytokine production (Figure 4a). When comparing NSCLC BM patients with healthy controls, we found significantly higher frequencies of IL-10 in CD4^+^ and CD8^+^ T-cells of BM patients (P= 0.003 and 0.015 respectively, Welch’s t-test) (Figure 4 b). In analogy to our finding of increased T_H_17 CD4^+^ T helper cells, we also could see a significant increase of IL-17 production in CD4^+^ T cells (P=0.006, Welch’s t-test). When looking at the cytokine production in NK cells, we see higher mean frequencies of NK cells expressing IL-2, IL-4, and IL-17 in NSCLC BM patients compared to healthy controls (Figure 4 b, P=0.004, 0.002 and 0.034 respectively, Welch’s t-test).

**Figure 4:**
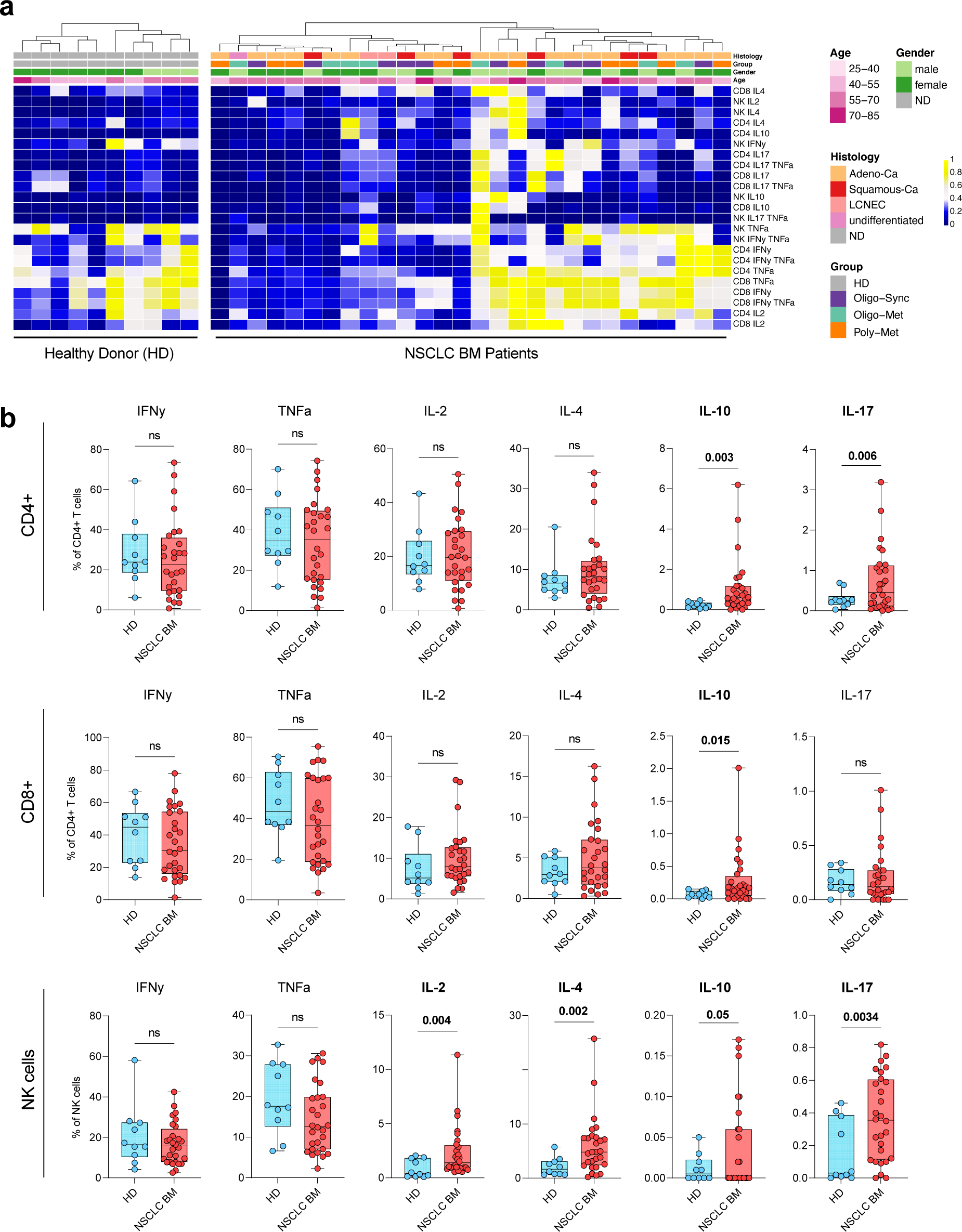
Flow cytometric T-cells cytokine analysis of stimulated peripheral blood lymphocytes. (a.) Heatmap representing the median expression of cytokines in NSCLC BM and HD samples. (b.) Box plots depicting expression of IFN***γ***, TNF***α***, IL-2, IL-4, IL-10, and IL-17 in CD4, CD8 T-cells, and NK cells. Mean ± SEM, t test. P values are defined as * < 0.05; ** < 0.01; and *** < 0.001.

On the other hand, no significant differences were observed in the T-cell metabolism profile (FACS panel 4) between NSCLC BM patients and HD (no data shown).

### Peripheral blood immune profile of patients with oligo-metastatic disease

When all analyzed patient samples were stratified according to their metastatic phenotype, the most intriguing findings were related to T-cell subtype differentiation. We assessed the frequencies of the different stages of T-cell development as follows: Naïve: (CD45RA+, CD27+, CD28+, CCR7+), Central memory: (CD45RA−, CD27+, CD28+, CCR7+), Early memory (Early: CD45RA-, CD27+, CD28+, CCR7-), Early-like memory (Early-like: CD45RA-, CD27-, CD28+, CCR7+), Intermediate (CD45RA-, CD27+, CD28-, CCR7-), Effector memory (effector memory cells: CD45RA−, CD27−, CD28+/−, CCR7−) and TeffRA+ (T effectors type RA+: CD45RA+, CD27−, CD28−, CCR7−) ^19^.

Supervised analyses revealed significant mean differences in the percentages of CD4^+^ T-cell populations between the three brain metastatic groups, whereas no such differences were observed in CD8^+^ T-cells (Figure 5a). These CD4^+^ subtypes, which reflect the composition of differentiation and activation in the peripheral blood compartment and play crucial roles in mediating adaptive immunity, exhibited fewer CD4^+^ effector type RA^+^ cells (T_Eff_RA^+^) among oligo-synchronous patients compared to oligo-metachronous (P=0.053, Wilcoxon test) and poly brain metastatic groups (P=0.020, Wilcoxon test), respectively (Figure 5b). Similarly, the median percentage of CD4^+^ effector type RA^-^ cells (T_Eff_RA^-^)/effector memory (Tem) cells were significantly lower in oligo-synchronous patients in comparison to the other two groups (P=0.021, and P=0.002 respectively, Wilcoxon test). In contrast, no significant differences were seen among the three brain metastatic groups in terms of CD4 early, early-like, intermediate, or central memory cells. Finally, the median percentage of CD4 naïve cells was significantly higher in oligo-synchronous (P=0.012, Wilcoxon test) as compared to oligo-metachronous (Figure 5b). To assess if pre-treatment had an impact on the T-cell functional differentiation profiles of BM groups, a comparison was made between untreated samples from oligo-synchronous and poly BM. The results of this analysis did not lead to an alteration of the significant differences observed. However, when only untreated oligo-synchronous samples were compared with pre-treated oligo-metachronous samples, no significant differences could be observed anymore (Supplemental Figure7). This confirms that treatment is not related to the differences seen between the BM groups.

**Figure 5:**
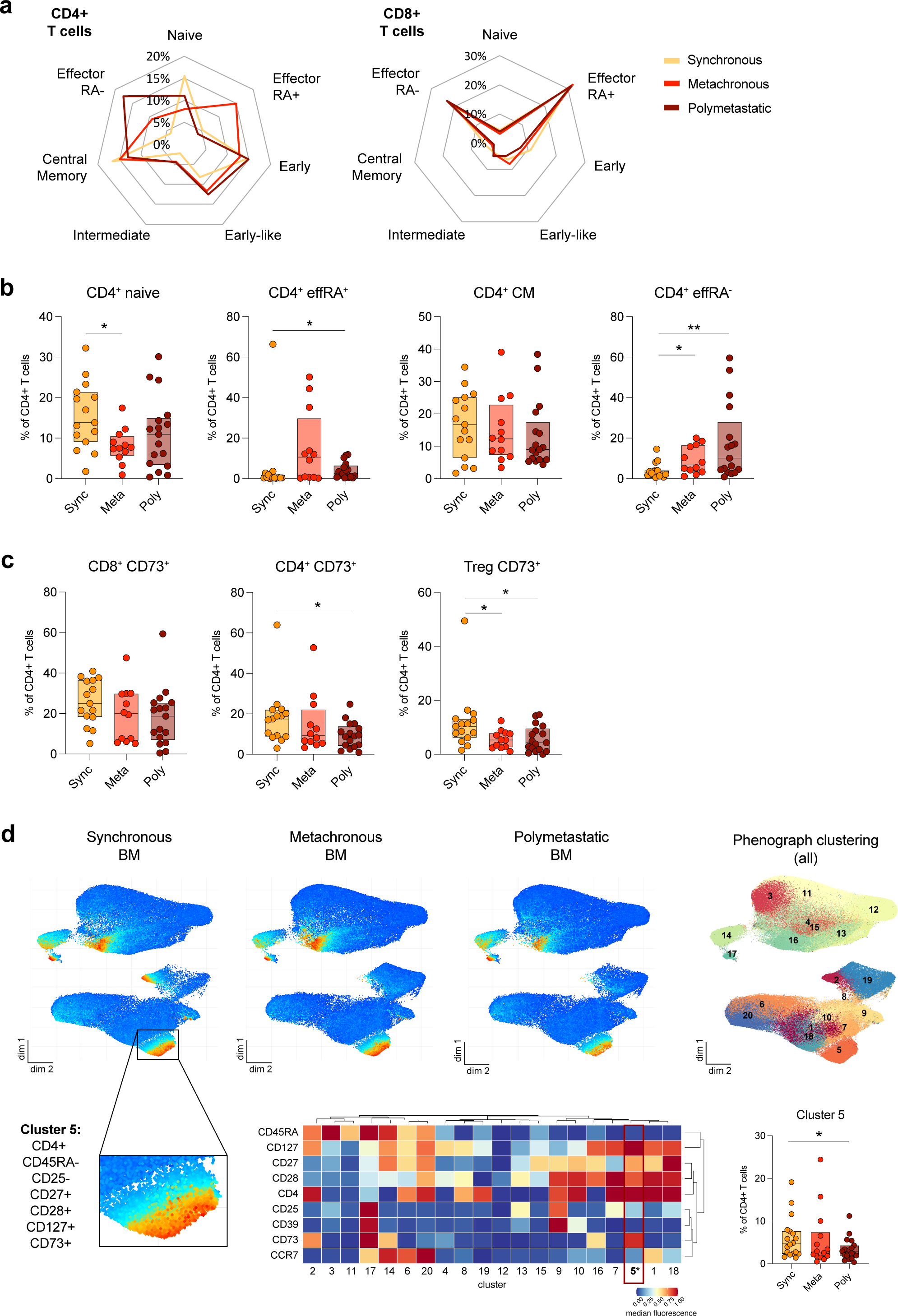
Flow cytometric immunophenotyping of T-cell differentiation. (a.) Spider plots of CD4+ and CD8+ differentiation phenotypes of peripheral blood in BM groups. Populations are defined by expressions of CD45RA, CD27, CD28, and CCR7 markers. Naïve: (CD45RA+, CD27+, CD28+, CCR7+), TeffRA+: T effectors type RA+ (CD45RA+, CD27−, CD28−, CCR7−), Early (CD45RA-, CD27+, CD28+, CCR7-), Early-like (CD45RA-, CD27-, CD28+, CCR7+), Intermediate (CD45RA-, CD27+, CD28-, CCR7-), Central Memory: (CD45RA−, CD27+, CD28+, CCR7+), and TeffRA-: effector memory cells (CD45RA−, CD27−, CD28−, CCR7− (b.) Box plots depicting expression of CD4+ naïve, TeffRA+, central memory, and TeffRA-expressions between BM groups. (c.) Box plots of CD73 expressions in CD8+, CD4+, and Treg cells between BM groups. (d.) Phonograph clustering of different markers in the three groups combined, and individual UMAPs display CD73 expression circled in red (cluster2) in Synchronous, Metachronous, and polymetastatic BM. Heatmap of markers expression in cluster 2: CD4+, CD45RA+, CD127+ and CD73+. Values describe population size as percentages. (Sync: oligo-synchronous BM, Meta: oligo-metachronous BM, and poly: poly metastasis).

Both the surface ectonucleotidases CD73 and CD39 which are activation/exhaustion markers, were assessed in the cancer patients using the same flow cytometry panel assessing T-cell differentiation. CD73 expression was investigated in both CD4^+^ and CD8^+^ T-cells and a significant mean difference was seen in CD4^+^ cells between the oligo-synchronous and the poly BM group (P=0.022, Wilcoxon test), however, no difference was observed in CD73 expression by CD8^+^ T-cells between any of the brain metastatic patient groups (Figure 5c). This observed difference in CD4^+^ was also significant for the subset of regulatory T-cells (Tregs) between the oligo-synchronous and poly BM groups (P=0.023, Wilcoxon test) and between the oligo-synchronous and -metachronous BM groups (P=0.016, Wilcoxon test) (Figure 5c). No difference in the percentage of any of the T-cell subgroups expressing CD39 could be discerned between the patient groups. Interestingly, also the unsupervised UMAP analysis indicated a specific role of CD73 in the oligo-synchronous group, shown by a difference in the percentage of CD73-expressing cells observed between the groups in cluster 5 and at the tip of cluster 2 (Figure 5d). The heat map shows that the cluster 5 mainly consisted of CD4^+^, CD27^+^, CD28^+^, CD127^+^, CD73^+^ and CD45RA^-^, CD25^-^ cells. These cells were significantly enriched in the oligo-synchronous group compared to the poly-metastatic group (P=0.044, t-test). These results indicate that the metastatic phenotype of oligo-synchronous metastasis is associated with an altered CD4^+^ T-cell differentiation.

## Discussion

Systemically, tumor cells and the peripheral immune system mutually influence each other, resulting in what is known as a systemic immune environment ^20^. Thus, we conducted a comprehensive analysis of both local and peripheral immunological characteristics of NSCLC patients with different brain metastatic patterns.

The immunophenotyping of the brain metastatic tissue, identified CD68^+^ cells to be the most prevalent immune cell type in both the peri- and the intra-tumoral regions. Furthermore, we detected significantly more CD68^+^ cells and lymphocytes (TILs) in the peri-tumoral regions compared to the intra-tumoral regions. However, similar to other studies we found that the number of TILs was noticeably lower than what has been reported for primary lung tumors, indicating the presence of a more immunosuppressive microenvironment in the brain ^21–23^.

To the best of our knowledge, no previous studies have explored the immune profiles among different brain metastatic patterns. Our analysis revealed a distinct augmentation of CD4^+^ T-cells among the oligo-synchronous BM patients. This difference was particularly noticeable in terms of intra-tumoral CD4^+^ T-cell infiltration. To further investigate the CD4^+^ T-cells, we analyzed their expression of FoxP3, which serves as a marker for Tregs. Surprisingly, although there was no significant difference in the percentage of T cells expressing FoxP3 among the different BM groups, we observed a positive correlation between FoxP3-expressing infiltrating cells in intra-tumoral regions and overall survival. Limited data are available regarding the prognostic role of FoxP3^+^ cells in BM ^24,25^. However, in line with our findings, another study focusing on NSCLC BM reported a positive correlation between OS and FoxP3 density within tumor lesions, but not in the peri-tumoral areas ^21^.

The multi-parametric immune phenotyping of peripheral blood identified several immune cell populations differentiating HD from BM patients. We identified significantly less CD8^+^ T-cells expressing KLRG1 in NSCLC patients BM compared to HD. KLRG1 serves as a marker for terminal differentiation and senescence ^26^. Furthermore, KLRG1 downregulation on CD8^+^ T-cells may indicate functional exhaustion and a potential impairment in the CD8-mediated antitumor immune response ^27,28^. These observations support the notion that the immune system’s ability to effectively target cancer cells in the brain may be compromised in BM NSCLC patients. Moreover, our investigation showed a significant increase in both CD4^+^ T_H_17 cells and elevated IL-17 cytokine production in the peripheral blood of BM patients compared to healthy individuals. The involvement of T_H_17 cells, characterized by production of IL-17, has been extensively studied in various cancer entities ^29^. While a pro-inflammatory phenotype of T_H_17 cells can contribute to promoting antitumor immunity, these cells can also exhibit pro-tumorigenic effects and immunosuppressive functions, likely related their plasticity, enabling them to adopt different phenotypes depending on the surrounding microenvironment ^30^. Previous studies have reported higher T_H_17 lymphocytes in peripheral blood of patients with NSCLC ^31,32^. Our data, supported by previous findings, thus indicate suggest an involvement of T_H_17-mediated responses in the immunopathology of NSCLC brain metastases. Similarly, we identified a CD4-related inverse relation between reduced TIGIT and increased T_H_17 expression, in BM patients indicating again a less immunosuppressive state compared to HD.

In our study, we furthermore observed a significant upregulation of IL-10 producing CD4^+^ and CD8^+^ T-cells in the BM patients, compared to healthy controls. This finding is consistent with the previously reported increase in IL-10 levels in the peripheral blood of cancer patients, in which higher levels also predicted a worse outcome ^33,34^. Notably, the elevation of IL-10 in our cohort of NSCLC patients with BM complements our earlier observations of an overall inflammatory phenotype in NSCLC, characterized by heightened levels of T_H_17 cells and IL-17. The concurrent upregulation of IL-10 and IL-17 underscores the interactive balance between pro- and anti-inflammatory responses in the immune system as modulated by cancer.

In summary, our study revealed significant alterations in the peripheral immune system, particularly within the CD4^+^ T-cell compartment, when comparing the global effects of NSCLC BM and healthy individuals.

The role of CD4^+^ T-cells in immuno-oncology has gained increasing recognition ^35^. Notably, CD4^+^ T-cells, including T-helpers and Tregs, can affect metastatic progression independently of CD8^+^ T-cells ^18,36,37^. Given the highly specialized immune environment of the brain, data on CD4^+^ T-cell composition in the brain tumor microenvironment is scarce. A recent single-cell multi-omics analyses of different brain metastases indicated, similarly to our findings, that CD4^+^ T-cells show much greater degree of phenotypic heterogeneity compared to CD8^+^ T-cells ^38^. Furthermore, the authors showed that CD8^+^ T-cells most commonly were effector memory cells, whereas CD4^+^ cells again displayed a more heterogeneous continuum across effector memory and central memory T-cells. Our findings, along with previous research, indicate a vital role of CD4^+^ T-cells in mediating antitumor immunity in NSCLC BM, irrespective of CD8^+^ T-cells.

When the patients were divided into the different metastatic phenotypes, no significant differences in CD4^+^ and CD8^+^ T-cell population size were noticed between the BM groups. However, similarly to the BM tissue results, the distribution of the CD4^+^ differentiation phenotypes was shifted. When we examined the expression patterns of different T-cell differentiation subtypes, our results showed remarkable differences in CD4^+^ subtypes patterns between BM groups, while again such distinction was not noticed in CD8^+^ subtypes. This might indicate that metastatic patterns trigger different immune responses specifically in the peripheral CD4^+^ compartment. We found more CD4^+^ naïve cells, whereas T_Eff_RA^+^ and T_Eff_RA^-^ cells were significantly less observed in the oligo-synchronous BM group. Naive T-cells act as immune surveillance circulating in the blood and respond to foreign antigens. As a result, they differentiate into effector cells T_Eff_RA^+^ that kill or help other immune cells to attack the pathogen or later to e.g. T_Eff_RA^-^ cells, which in turn perform the memory immune response when the infection reoccurs ^39,40^. Our results thus indicate that patients with oligo-synchronous BM have a less activated immune system indicating that a brain metastasis alone might not evoke a strong immune response in the periphery, nonetheless allowing the outgrowth of a tumor in the brain.

Furthermore, our study showed in general an upregulation of CD73 on especially CD4^+^ T-cells in all BM patients compared to HD, with the strongest up-regulation among oligo-synchronous BM patients. The unsupervised analysis revealed a distinct cell population in oligo-synchronous BM, comprised of CD4^+^, CD27^+^, CD28^+^, CD127^+^, CD73^+^ and CD25^-^-cells. These cells are likely memory CD4^+^ cells. Notably, Doherty and colleagues identified effector-memory CD73^+^ CD4^+^ T-cells as T_H_17 cells ^41^, and Gourdin et al. demonstrated that CD73 identifies a subset of CD4^+^ effector T-cells enriched with T_H_17 populations ^42^. These findings align with the higher T_H_17 frequency observed in our BM cohort.

Beyond its well-described enzymatic function ^43^, CD73 functions as a lymphocyte differentiation antigen, implying involvement in lymphocyte maturation, development, and T-cell activation. CD73 also serves as an adhesion molecule, facilitating the binding of lymphocytes to the endothelium ^44–46^. While research on CD73 in the peripheral immune system is limited, its impact on tumor progression and anti-tumor responses in NSCLC has been observed in the TME ^47,48^. Given the multifaceted roles and mechanisms of CD73, careful interpretation of our results is warranted. Although the precise function of CD73 in BM-NSCLC necessitates further investigation, our data indicate intriguing differences among the different BM cohorts.

In summary, it has been proposed that peripheral CD4^+^ T-cell differentiation patterns can independently predict tumor progression in NSCLC ^49^. Our data further proves that this application also provides valuable information on the type of metastatic spread of BM patients. Although our cohort size was limited, we comprehensively analyzed the peripheral and microenvironment immune landscape of NSCLC BM patients. We focused here on T-cells and NKs, however, other immune cells most likely influence on BM formation. Furthermore, acomparison with non-BM NSCLC patients would be desirable, however, given the heterogeneous metastatic patterns in such cases and the mostly missing information about micrometastases in the brain in asymptomatic patients, impedes such comparative analyses.

Moreover, future studies need to define the biological role of these identified differences especially in driving oligo-metastatic disease.

In conclusion, our data show that NSCLC BM patients have a skewed systemic T- and NK cell phenotype compared to healthy individuals, with a surface marker profile that is indicative of dysregulation in both the CD4^+^ and CD8^+^ T cell compartments defined by KRLG1 and the T_H_17/IL-17 axis. In addition, a specific immune profile was identified separating especially the oligo-synchronous BM from the other groups in both peripheral blood as well as in brain tumor microenvironment. Specific CD4^+^ T-cell populations were altered in both the BM microenvironment of oligo-synchronous compared to the other BM groups as well as in the peripheral blood.

## Funding

This study was funded by the Wilhelm Sander-Stiftung (2017.075.1 SS, HW, MM) and Fritz Bender Stiftung (SJ, HW, MM) and Erich und Gertrud Roggenbuck-Stiftung (MM).

## Conflict of Interest

We declare no conflict of interest related to this manuscript.

## Authorship

MA, CLM, SS, GE, and JK performed the FACS experiments. MA, JM, MG, and HW performed and analyzed the IHC data. AP, SP, BA, JG, MW, KL and MM collected samples and annotated the clinical data. MA, LG, MR, AR, ET, SAJ, MM and HW analyzed the data. MA, SAJ, MM, and HW participated in drafting the manuscript. All authors have read and agreed to the published version of the manuscript.

## Acknowledgments

The authors would like to thank Sarina Heinemann, Katharina Kolbe, Svenja Zapf, and Jessica Stanik for excellent technical assistance. We would like to thank the UKE FACS Core Facility for providing support for the FACS analyses.

## Notes

### Competing Interest Statement

The authors have declared no competing interest.

## References

1. Sung H, Ferlay J, Siegel RL, et al. Global Cancer Statistics 2020: GLOBOCAN Estimates of Incidence and Mortality Worldwide for 36 Cancers in 185 Countries. CA Cancer J Clin. 2021; 71(3):209–249.

2. Villano JL, Durbin EB, Normandeau C, Thakkar JP, Moirangthem V, Davis FG. Incidence of brain metastasis at initial presentation of lung cancer. Neuro Oncol. 2015; 17(1):122–128.

3. Massague J, Ganesh K. Metastasis-Initiating Cells and Ecosystems. Cancer Discov. 2021; 11(4):971–994.

4. Piffko A, Asey B, Duhrsen L, et al. Clinical determinants impacting overall survival of patients with operable brain metastases from non-small cell lung cancer. Front Oncol. 2022; 12:951805.

5. Weichselbaum RR, Hellman S. Oligometastases revisited. Nat Rev Clin Oncol. 2011; 8(6):378–382.

6. Lussier YA, Khodarev NN, Regan K, et al. Oligo- and polymetastatic progression in lung metastasis(es) patients is associated with specific microRNAs. PLoS One. 2012; 7(12):e50141.

7. Gomez DR, Tang C, Zhang J, et al. Local Consolidative Therapy Vs. Maintenance Therapy or Observation for Patients With Oligometastatic Non-Small-Cell Lung Cancer: Long-Term Results of a Multi-Institutional, Phase II, Randomized Study. J Clin Oncol. 2019; 37(18):1558–1565.

8. Petty WJ, Urbanic JJ, Ahmed T, et al. Long-Term Outcomes of a Phase 2 Trial of Chemotherapy With Consolidative Radiation Therapy for Oligometastatic Non-Small Cell Lung Cancer. International journal of radiation oncology, biology, physics. 2018; 102(3):527–535.

9. Tsai CJ, Yang JT, Shaverdian N, et al. Standard-of-care systemic therapy with or without stereotactic body radiotherapy in patients with oligoprogressive breast cancer or non-small-cell lung cancer (Consolidative Use of Radiotherapy to Block [CURB] oligoprogression): an open-label, randomised, controlled, phase 2 study. Lancet. 2024; 403(10422):171–182.

10. Hanahan D, Weinberg RA. Hallmarks of cancer: the next generation. Cell. 2011; 144(5):646–674.

11. Kim R, Keam B, Kim S, et al. Differences in tumor microenvironments between primary lung tumors and brain metastases in lung cancer patients: therapeutic implications for immune checkpoint inhibitors. BMC Cancer. 2019; 19(1):19.

12. Karimi E, Yu MW, Maritan SM, et al. Single-cell spatial immune landscapes of primary and metastatic brain tumours. Nature. 2023; 614(7948):555–563.

13. Camy F, Karpathiou G, Dumollard JM, et al. Brain metastasis PD-L1 and CD8 expression is dependent on primary tumor type and its PD-L1 and CD8 status. J Immunother Cancer. 2020; 8(2).

14. Riebensahm C, Joosse SA, Mohme M, et al. Clonality of circulating tumor cells in breast cancer brain metastasis patients. Breast Cancer Res. 2019; 21(1):101.

15. Greenberg SA, Kong SW, Thompson E, Gulla SV. Co-inhibitory T cell receptor KLRG1: human cancer expression and efficacy of neutralization in murine cancer models. Oncotarget. 2019; 10(14):1399.

16. Joller N, Lozano E, Burkett PR, et al. Treg cells expressing the coinhibitory molecule TIGIT selectively inhibit proinflammatory Th1 and Th17 cell responses. Immunity. 2014; 40(4):569–581.

17. Bailey SR, Nelson MH, Himes RA, Li Z, Mehrotra S, Paulos CM. Th17 cells in cancer: the ultimate identity crisis. Frontiers in immunology. 2014; 5:276.

18. Speiser DE, Chijioke O, Schaeuble K, Münz C. CD4+ T cells in cancer. Nature Cancer. 2023:1–13.

19. Cossarizza A, Chang HD, Radbruch A, et al. Guidelines for the use of flow cytometry and cell sorting in immunological studies. Eur J Immunol. 2017; 47(10):1584–1797.

20. Li YD, Lamano JB, Lamano JB, et al. Tumor-induced peripheral immunosuppression promotes brain metastasis in patients with non-small cell lung cancer. Cancer Immunol Immunother. 2019; 68(9):1501–1513.

21. Ikarashi D, Okimoto T, Shukuya T, et al. Comparison of Tumor Microenvironments Between Primary Tumors and Brain Metastases in Patients With NSCLC. JTO Clin Res Rep. 2021; 2(10):100230.

22. Song Z, Yang L, Zhou Z, et al. Genomic profiles and tumor immune microenvironment of primary lung carcinoma and brain oligo-metastasis. Cell Death & Disease. 2021; 12(1):106.

23. Mansfield A, Aubry M, Moser J, et al. Temporal and spatial discordance of programmed cell death-ligand 1 expression and lymphocyte tumor infiltration between paired primary lesions and brain metastases in lung cancer. Annals of Oncology. 2016; 27(10):1953–1958.

24. Berghoff AS, Fuchs E, Ricken G, et al. Density of tumor-infiltrating lymphocytes correlates with extent of brain edema and overall survival time in patients with brain metastases. Oncoimmunology. 2016; 5(1):e1057388.

25. Harter PN, Bernatz S, Scholz A, et al. Distribution and prognostic relevance of tumor-infiltrating lymphocytes (TILs) and PD-1/PD-L1 immune checkpoints in human brain metastases. Oncotarget. 2015; 6(38):40836–40849.

26. Legat A, Speiser DE, Pircher H, Zehn D, Fuertes Marraco SA. Inhibitory receptor expression depends more dominantly on differentiation and activation than “exhaustion” of human CD8 T cells. Frontiers in immunology. 2013; 4:455.

27. Wherry EJ, Kurachi M. Molecular and cellular insights into T cell exhaustion. Nature Reviews Immunology. 2015; 15(8):486–499.

28. Herndler-Brandstetter D, Ishigame H, Shinnakasu R, et al. KLRG1+ effector CD8+ T cells lose KLRG1, differentiate into all memory T cell lineages, and convey enhanced protective immunity. Immunity. 2018; 48(4):716–729. e718.

29. Zou W, Restifo NP. TH17 cells in tumour immunity and immunotherapy. Nature Reviews Immunology. 2010; 10(4):248–256.

30. Karpisheh V, Ahmadi M, Abbaszadeh-Goudarzi K, et al. The role of Th17 cells in the pathogenesis and treatment of breast cancer. Cancer Cell International. 2022; 22(1):108.

31. Chen G, Zhang P-G, Li J-S, et al. Th17 cell frequency and IL-17A production in peripheral blood of patients with non-small-cell lung cancer. Journal of International Medical Research. 2020; 48(6):0300060520925948.

32. Song L, Ma S, Chen L, Miao L, Tao M, Liu H. LonglJterm prognostic significance of interleukinlJ17lJproducing T cells in patients with nonlJsmall cell lung cancer. Cancer science. 2019; 110(7):2100–2109.

33. Zhao S, Wu D, Wu P, Wang Z, Huang J. Serum IL-10 predicts worse outcome in cancer patients: a meta-analysis. PloS one. 2015; 10(10):e0139598.

34. Sung W-W, Wang Y-C, Lin P-L, et al. IL-10 promotes tumor aggressiveness via upregulation of CIP2A transcription in lung adenocarcinoma. Clinical Cancer Research. 2013; 19(15):4092–4103.

35. Miggelbrink AM, Jackson JD, Lorrey SJ, et al. CD4 T-cell exhaustion: does it exist and what are its roles in cancer? Clinical cancer research: an official journal of the American Association for Cancer Research. 2021; 27(21):5742.

36. DeNardo DG, Barreto JB, Andreu P, et al. CD4(+) T cells regulate pulmonary metastasis of mammary carcinomas by enhancing protumor properties of macrophages. Cancer Cell. 2009; 16(2):91–102.

37. Kruse B, Buzzai AC, Shridhar N, et al. CD4+ T cell-induced inflammatory cell death controls immune-evasive tumours. Nature. 2023:1–8.

38. Gonzalez H, Mei W, Robles I, et al. Cellular architecture of human brain metastases. Cell. 2022; 185(4):729–745. e720.

39. Yamaguchi K, Mishima K, Ohmura H, et al. Activation of central/effector memory T cells and T-helper 1 polarization in malignant melanoma patients treated with anti-programmed death-1 antibody. Cancer Sci. 2018; 109(10):3032–3042.

40. Saxena A, Dagur PK, Biancotto A. Multiparametric Flow Cytometry Analysis of Naive, Memory, and Effector T Cells. Methods Mol Biol. 2019; 2032:129–140.

41. Doherty GA, Bai A, Hanidziar D, et al. CD 73 is a phenotypic marker of effector memory T h17 cells in inflammatory bowel disease. European journal of immunology. 2012; 42(11):3062–3072.

42. Gourdin N, Bossennec M, Rodriguez C, et al. Autocrine adenosine regulates tumor polyfunctional CD73+ CD4+ effector T cells devoid of immune checkpoints. Cancer research. 2018; 78(13):3604–3618.

43. Borsellino G, Kleinewietfeld M, Di Mitri D, et al. Expression of ectonucleotidase CD39 by Foxp3+ Treg cells: hydrolysis of extracellular ATP and immune suppression. Blood. 2007; 110(4):1225–1232.

44. Colgan SP, Eltzschig HK, Eckle T, Thompson LF. Physiological roles for ecto-5’-nucleotidase (CD73). Purinergic Signal. 2006; 2(2):351–360.

45. Resta R, Yamashita Y, Thompson LF. Ecto-enzyme and signaling functions of lymphocyte CD73. Immunol Rev. 1998; 161:95–109.

46. Yamashita Y, Hooker SW, Jiang H, et al. CD73 expression and fyn-dependent signaling on murine lymphocytes. Eur J Immunol. 1998; 28(10):2981–2990.

47. Tu E, McGlinchey K, Wang J, et al. Anti-PD-L1 and anti-CD73 combination therapy promotes T cell response to EGFR-mutated NSCLC. JCI Insight. 2022; 7(3).

48. Inoue Y, Yoshimura K, Kurabe N, et al. Prognostic impact of CD73 and A2A adenosine receptor expression in non-small-cell lung cancer. Oncotarget. 2017; 8(5):8738–8751.

49. Yang P, Ma J, Yang X, Li W. Peripheral CD4+ naive/memory ratio is an independent predictor of survival in non-small cell lung cancer. Oncotarget. 2017; 8(48):83650–83659.

