## Supplemental tables and figures for "NSCLC patients with oligo-metastatic brain disease show an altered CD4 T-cells immune profile"

### Slide 1
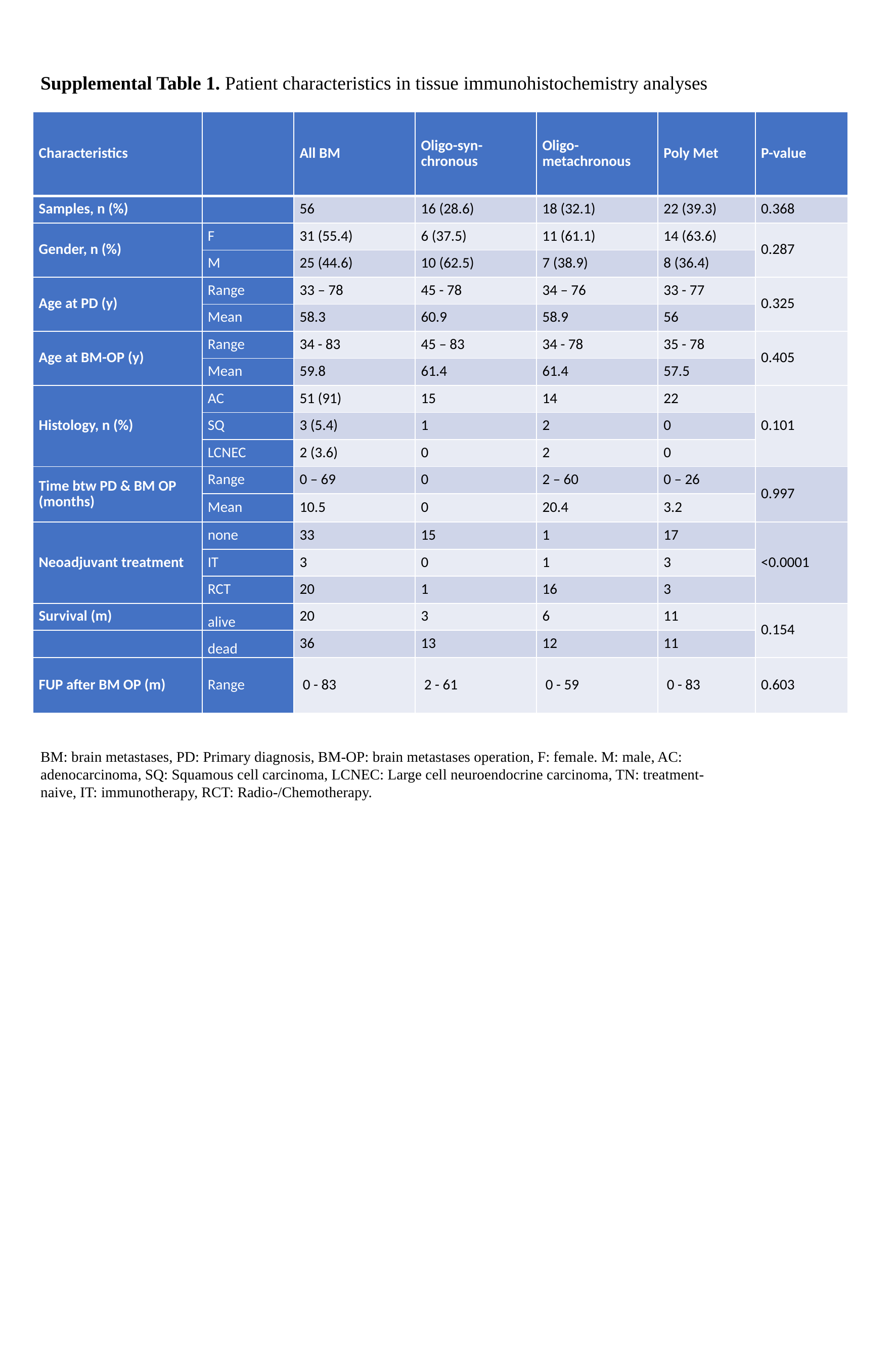

Supplemental Table 1. Patient characteristics in tissue immunohistochemistry analyses
| Characteristics | | All BM | Oligo-syn-chronous | Oligo-metachronous | Poly Met | P-value |
| --- | --- | --- | --- | --- | --- | --- |
| Samples, n (%) | | 56 | 16 (28.6) | 18 (32.1) | 22 (39.3) | 0.368 |
| Gender, n (%) | F | 31 (55.4) | 6 (37.5) | 11 (61.1) | 14 (63.6) | 0.287 |
| | M | 25 (44.6) | 10 (62.5) | 7 (38.9) | 8 (36.4) | |
| Age at PD (y) | Range | 33 – 78 | 45 - 78 | 34 – 76 | 33 - 77 | 0.325 |
| | Mean | 58.3 | 60.9 | 58.9 | 56 | |
| Age at BM-OP (y) | Range | 34 - 83 | 45 – 83 | 34 - 78 | 35 - 78 | 0.405 |
| | Mean | 59.8 | 61.4 | 61.4 | 57.5 | |
| Histology, n (%) | AC | 51 (91) | 15 | 14 | 22 | 0.101 |
| | SQ | 3 (5.4) | 1 | 2 | 0 | |
| | LCNEC | 2 (3.6) | 0 | 2 | 0 | |
| Time btw PD & BM OP (months) | Range | 0 – 69 | 0 | 2 – 60 | 0 – 26 | 0.997 |
| | Mean | 10.5 | 0 | 20.4 | 3.2 | |
| Neoadjuvant treatment | none | 33 | 15 | 1 | 17 | <0.0001 |
| | IT | 3 | 0 | 1 | 3 | |
| | RCT | 20 | 1 | 16 | 3 | |
| Survival (m) | alive | 20 | 3 | 6 | 11 | 0.154 |
| | dead | 36 | 13 | 12 | 11 | |
| FUP after BM OP (m) | Range | 0 - 83 | 2 - 61 | 0 - 59 | 0 - 83 | 0.603 |
BM: brain metastases, PD: Primary diagnosis, BM-OP: brain metastases operation, F: female. M: male, AC: adenocarcinoma, SQ: Squamous cell carcinoma, LCNEC: Large cell neuroendocrine carcinoma, TN: treatment-naive, IT: immunotherapy, RCT: Radio-/Chemotherapy.

### Slide 2
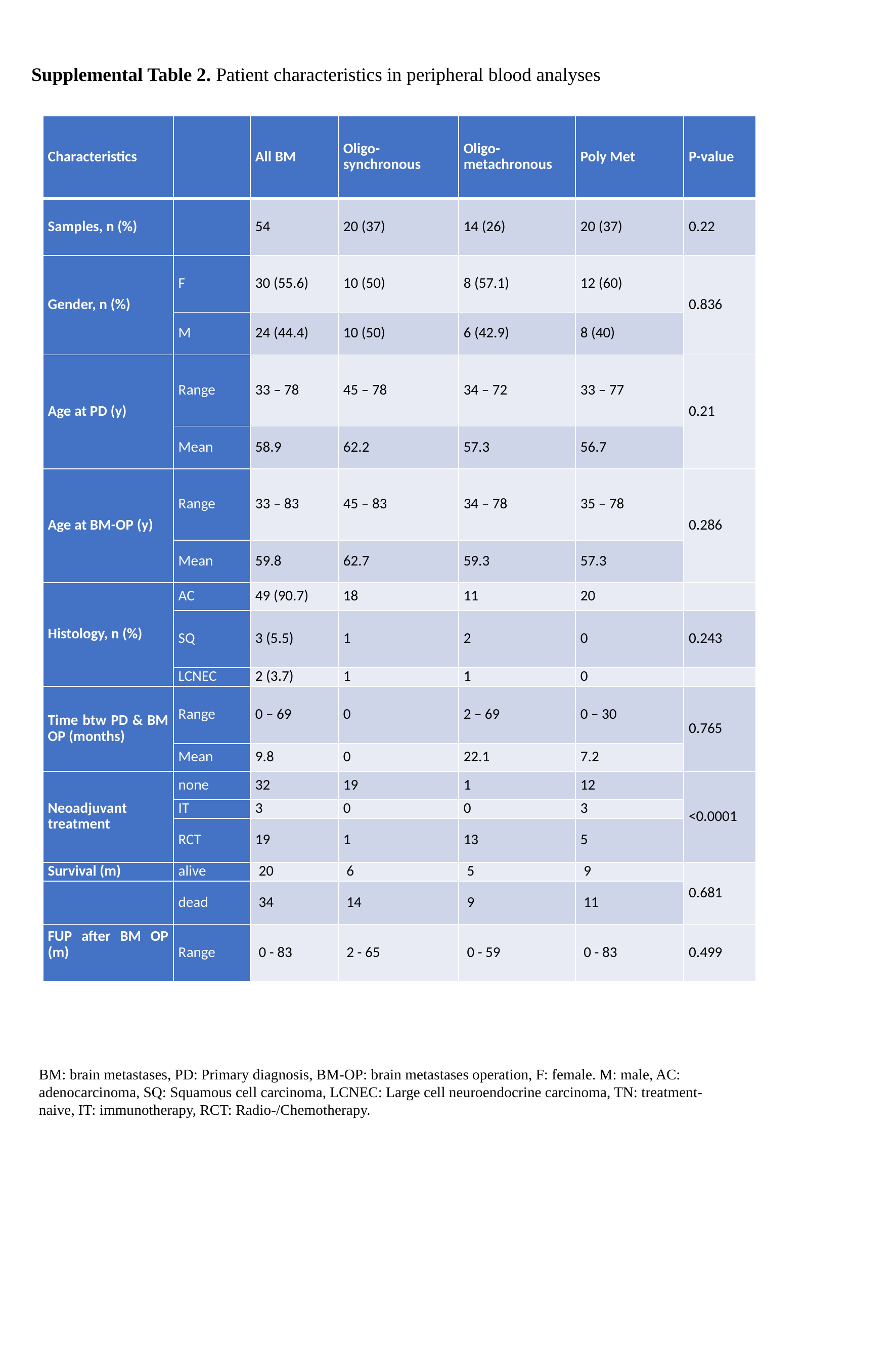

Supplemental Table 2. Patient characteristics in peripheral blood analyses
| Characteristics | | All BM | Oligo-synchronous | Oligo-metachronous | Poly Met | P-value |
| --- | --- | --- | --- | --- | --- | --- |
| Samples, n (%) | | 54 | 20 (37) | 14 (26) | 20 (37) | 0.22 |
| Gender, n (%) | F | 30 (55.6) | 10 (50) | 8 (57.1) | 12 (60) | 0.836 |
| | M | 24 (44.4) | 10 (50) | 6 (42.9) | 8 (40) | |
| Age at PD (y) | Range | 33 – 78 | 45 – 78 | 34 – 72 | 33 – 77 | 0.21 |
| | Mean | 58.9 | 62.2 | 57.3 | 56.7 | |
| Age at BM-OP (y) | Range | 33 – 83 | 45 – 83 | 34 – 78 | 35 – 78 | 0.286 |
| | Mean | 59.8 | 62.7 | 59.3 | 57.3 | |
| Histology, n (%) | AC | 49 (90.7) | 18 | 11 | 20 | |
| | SQ | 3 (5.5) | 1 | 2 | 0 | 0.243 |
| | LCNEC | 2 (3.7) | 1 | 1 | 0 | |
| Time btw PD & BM OP (months) | Range | 0 – 69 | 0 | 2 – 69 | 0 – 30 | 0.765 |
| | Mean | 9.8 | 0 | 22.1 | 7.2 | |
| Neoadjuvant treatment | none | 32 | 19 | 1 | 12 | <0.0001 |
| | IT | 3 | 0 | 0 | 3 | |
| | RCT | 19 | 1 | 13 | 5 | |
| Survival (m) | alive | 20 | 6 | 5 | 9 | 0.681 |
| | dead | 34 | 14 | 9 | 11 | |
| FUP after BM OP (m) | Range | 0 - 83 | 2 - 65 | 0 - 59 | 0 - 83 | 0.499 |
BM: brain metastases, PD: Primary diagnosis, BM-OP: brain metastases operation, F: female. M: male, AC: adenocarcinoma, SQ: Squamous cell carcinoma, LCNEC: Large cell neuroendocrine carcinoma, TN: treatment-naive, IT: immunotherapy, RCT: Radio-/Chemotherapy.

### Slide 3
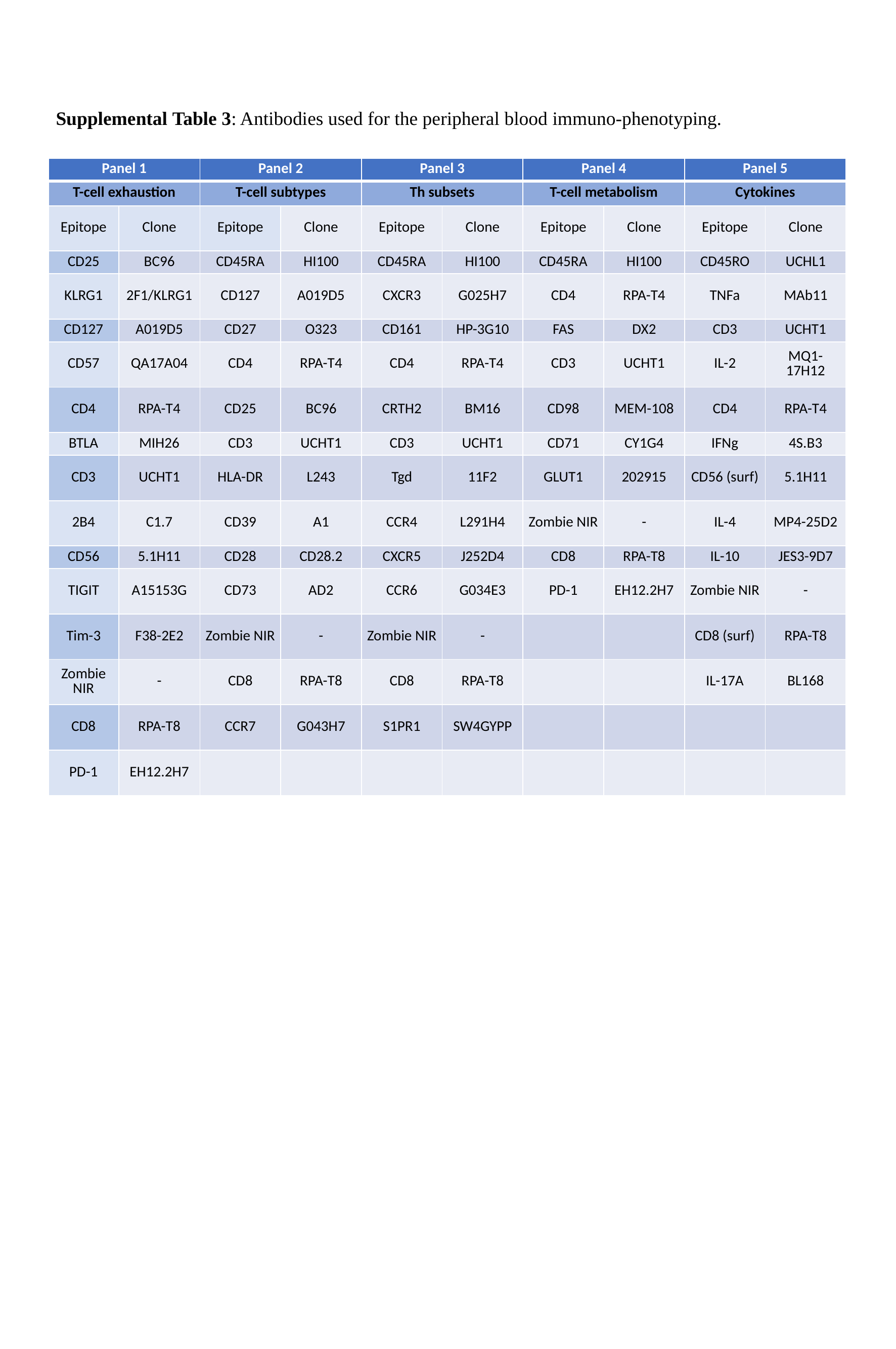

Supplemental Table 3: Antibodies used for the peripheral blood immuno-phenotyping.
| Panel 1 | | Panel 2 | | Panel 3 | | Panel 4 | | Panel 5 | |
| --- | --- | --- | --- | --- | --- | --- | --- | --- | --- |
| T-cell exhaustion | | T-cell subtypes | | Th subsets | | T-cell metabolism | | Cytokines | |
| Epitope | Clone | Epitope | Clone | Epitope | Clone | Epitope | Clone | Epitope | Clone |
| CD25 | BC96 | CD45RA | HI100 | CD45RA | HI100 | CD45RA | HI100 | CD45RO | UCHL1 |
| KLRG1 | 2F1/KLRG1 | CD127 | A019D5 | CXCR3 | G025H7 | CD4 | RPA-T4 | TNFa | MAb11 |
| CD127 | A019D5 | CD27 | O323 | CD161 | HP-3G10 | FAS | DX2 | CD3 | UCHT1 |
| CD57 | QA17A04 | CD4 | RPA-T4 | CD4 | RPA-T4 | CD3 | UCHT1 | IL-2 | MQ1-17H12 |
| CD4 | RPA-T4 | CD25 | BC96 | CRTH2 | BM16 | CD98 | MEM-108 | CD4 | RPA-T4 |
| BTLA | MIH26 | CD3 | UCHT1 | CD3 | UCHT1 | CD71 | CY1G4 | IFNg | 4S.B3 |
| CD3 | UCHT1 | HLA-DR | L243 | Tgd | 11F2 | GLUT1 | 202915 | CD56 (surf) | 5.1H11 |
| 2B4 | C1.7 | CD39 | A1 | CCR4 | L291H4 | Zombie NIR | - | IL-4 | MP4-25D2 |
| CD56 | 5.1H11 | CD28 | CD28.2 | CXCR5 | J252D4 | CD8 | RPA-T8 | IL-10 | JES3-9D7 |
| TIGIT | A15153G | CD73 | AD2 | CCR6 | G034E3 | PD-1 | EH12.2H7 | Zombie NIR | - |
| Tim-3 | F38-2E2 | Zombie NIR | - | Zombie NIR | - | | | CD8 (surf) | RPA-T8 |
| Zombie NIR | - | CD8 | RPA-T8 | CD8 | RPA-T8 | | | IL-17A | BL168 |
| CD8 | RPA-T8 | CCR7 | G043H7 | S1PR1 | SW4GYPP | | | | |
| PD-1 | EH12.2H7 | | | | | | | | |

### Slide 4
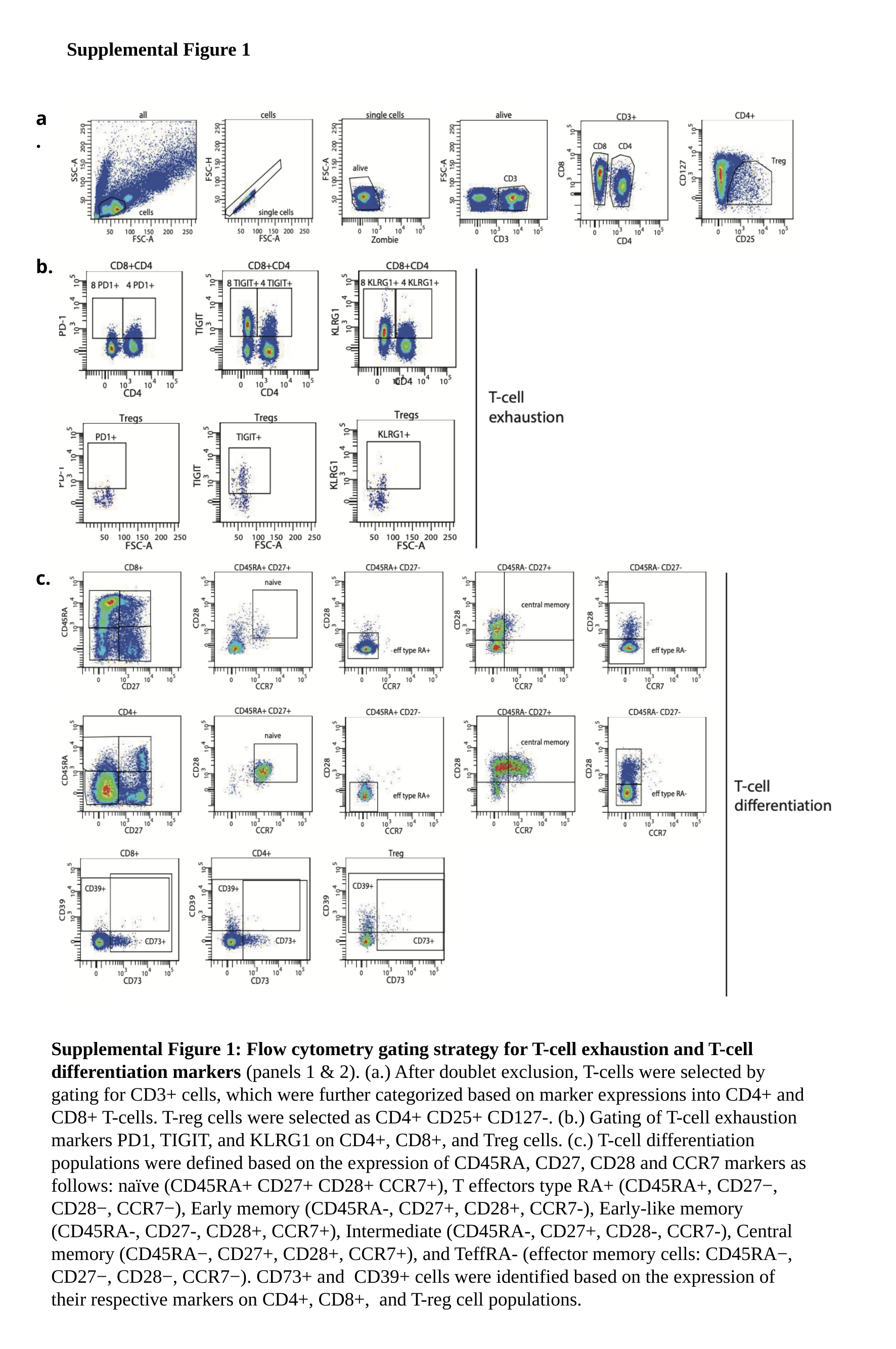

Supplemental Figure 1
a.
b.
c.
Supplemental Figure 1: Flow cytometry gating strategy for T-cell exhaustion and T-cell differentiation markers (panels 1 & 2). (a.) After doublet exclusion, T-cells were selected by gating for CD3+ cells, which were further categorized based on marker expressions into CD4+ and CD8+ T-cells. T-reg cells were selected as CD4+ CD25+ CD127-. (b.) Gating of T-cell exhaustion markers PD1, TIGIT, and KLRG1 on CD4+, CD8+, and Treg cells. (c.) T-cell differentiation populations were defined based on the expression of CD45RA, CD27, CD28 and CCR7 markers as follows: naïve (CD45RA+ CD27+ CD28+ CCR7+), T effectors type RA+ (CD45RA+, CD27−, CD28−, CCR7−), Early memory (CD45RA-, CD27+, CD28+, CCR7-), Early-like memory (CD45RA-, CD27-, CD28+, CCR7+), Intermediate (CD45RA-, CD27+, CD28-, CCR7-), Central memory (CD45RA−, CD27+, CD28+, CCR7+), and TeffRA- (effector memory cells: CD45RA−, CD27−, CD28−, CCR7−). CD73+ and CD39+ cells were identified based on the expression of their respective markers on CD4+, CD8+, and T-reg cell populations.

### Slide 5
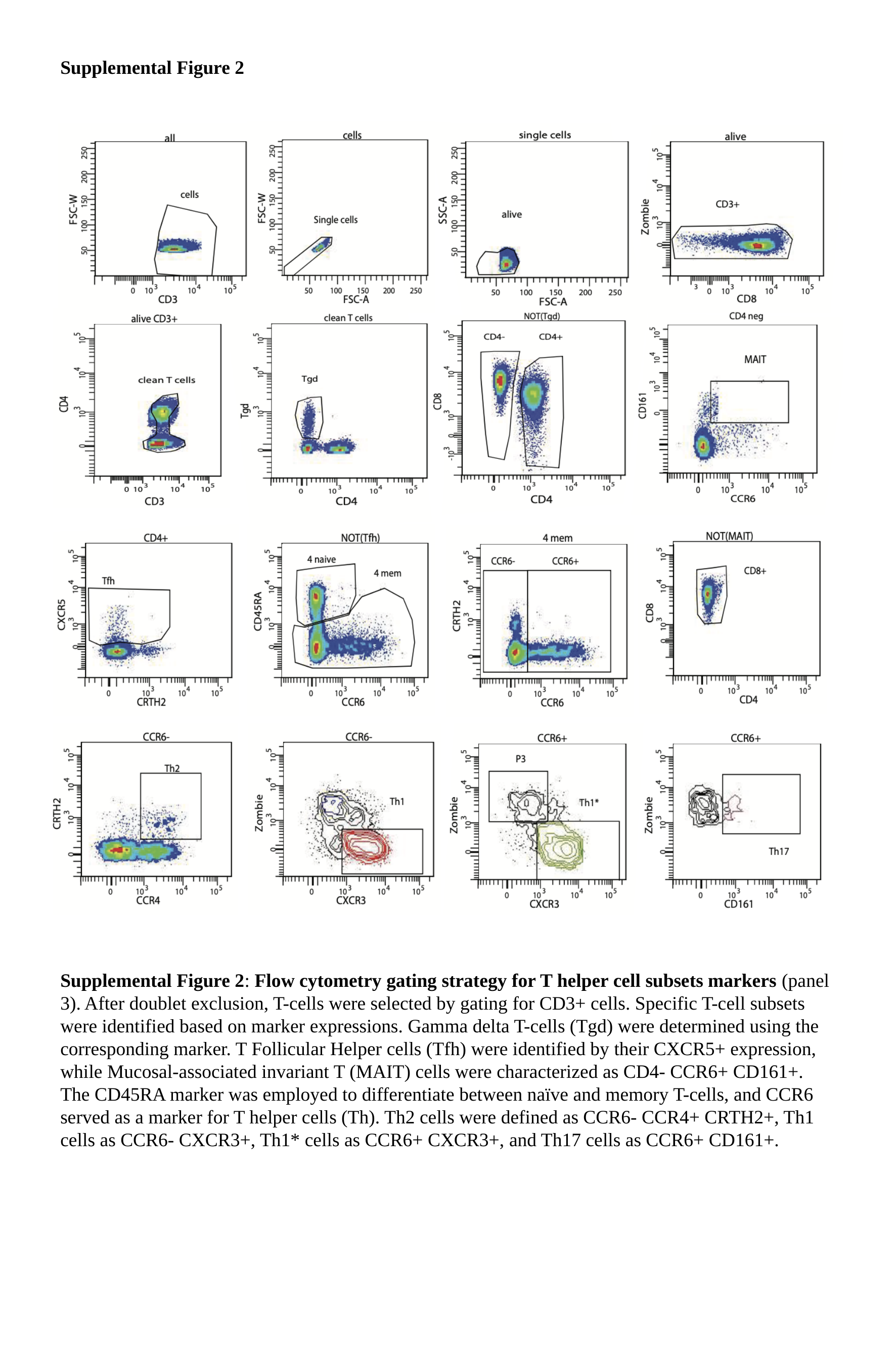

Supplemental Figure 2
Supplemental Figure 2: Flow cytometry gating strategy for T helper cell subsets markers (panel 3). After doublet exclusion, T-cells were selected by gating for CD3+ cells. Specific T-cell subsets were identified based on marker expressions. Gamma delta T-cells (Tgd) were determined using the corresponding marker. T Follicular Helper cells (Tfh) were identified by their CXCR5+ expression, while Mucosal-associated invariant T (MAIT) cells were characterized as CD4- CCR6+ CD161+. The CD45RA marker was employed to differentiate between naïve and memory T-cells, and CCR6 served as a marker for T helper cells (Th). Th2 cells were defined as CCR6- CCR4+ CRTH2+, Th1 cells as CCR6- CXCR3+, Th1* cells as CCR6+ CXCR3+, and Th17 cells as CCR6+ CD161+.

### Slide 6
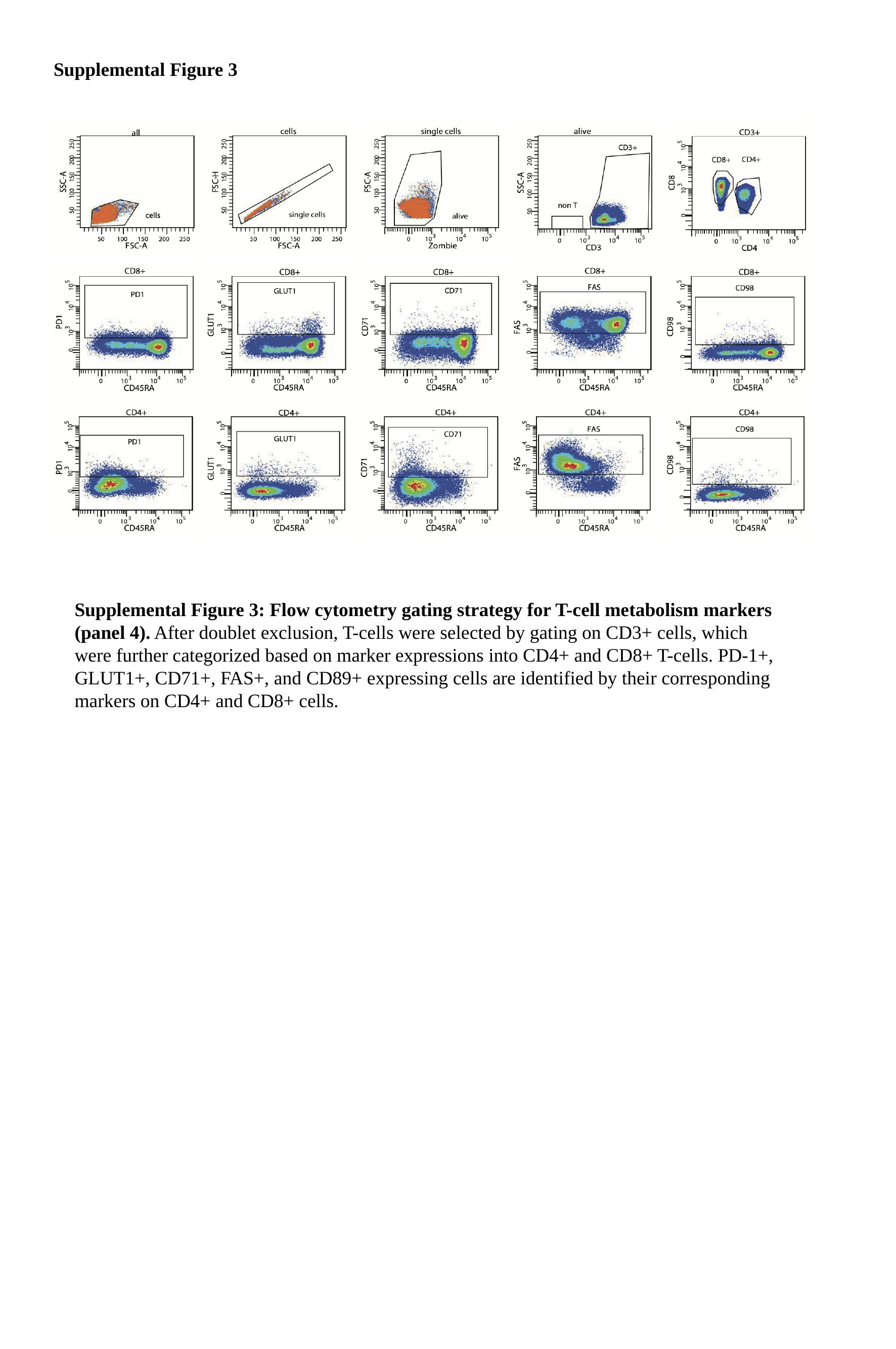

Supplemental Figure 3
Supplemental Figure 3: Flow cytometry gating strategy for T-cell metabolism markers (panel 4). After doublet exclusion, T-cells were selected by gating on CD3+ cells, which were further categorized based on marker expressions into CD4+ and CD8+ T-cells. PD-1+, GLUT1+, CD71+, FAS+, and CD89+ expressing cells are identified by their corresponding markers on CD4+ and CD8+ cells.

### Slide 7
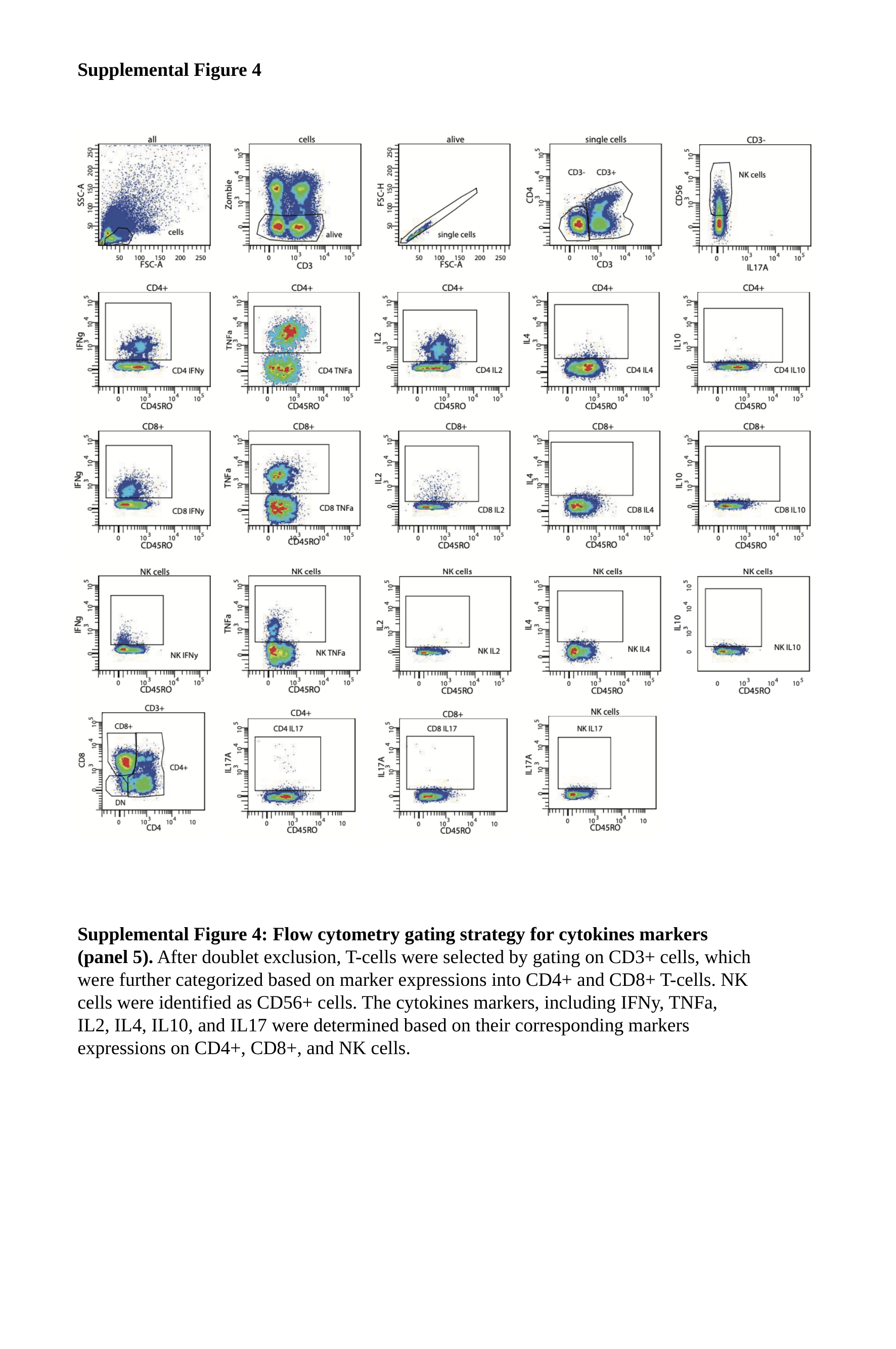

Supplemental Figure 4
Supplemental Figure 4: Flow cytometry gating strategy for cytokines markers (panel 5). After doublet exclusion, T-cells were selected by gating on CD3+ cells, which were further categorized based on marker expressions into CD4+ and CD8+ T-cells. NK cells were identified as CD56+ cells. The cytokines markers, including IFNy, TNFa, IL2, IL4, IL10, and IL17 were determined based on their corresponding markers expressions on CD4+, CD8+, and NK cells.

### Slide 8
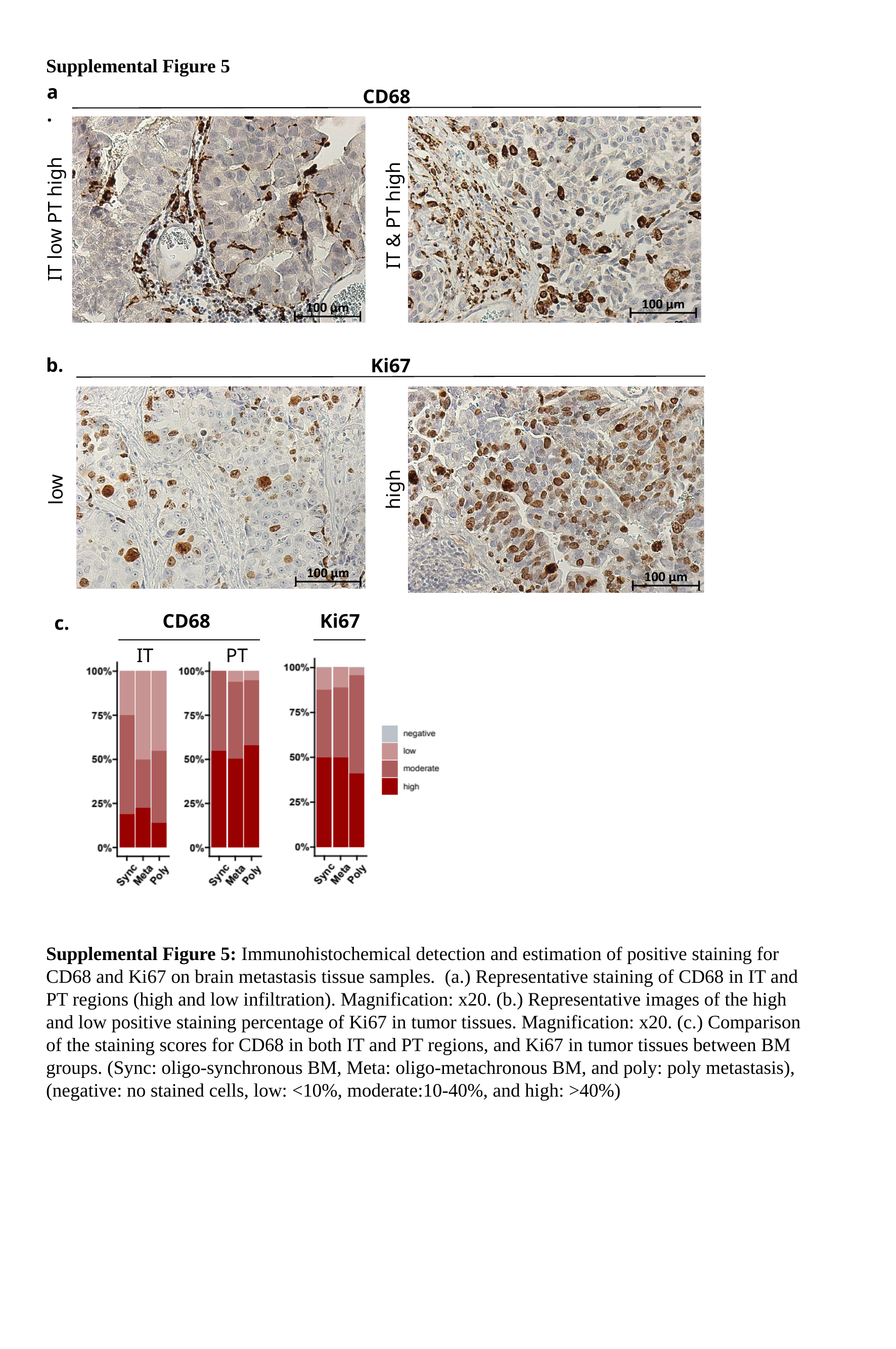

Supplemental Figure 5
a.
CD68
IT & PT high
IT low PT high
b.
Ki67
high
low
CD68
IT
PT
Ki67
c.
Supplemental Figure 5: Immunohistochemical detection and estimation of positive staining for CD68 and Ki67 on brain metastasis tissue samples. (a.) Representative staining of CD68 in IT and PT regions (high and low infiltration). Magnification: x20. (b.) Representative images of the high and low positive staining percentage of Ki67 in tumor tissues. Magnification: x20. (c.) Comparison of the staining scores for CD68 in both IT and PT regions, and Ki67 in tumor tissues between BM groups. (Sync: oligo-synchronous BM, Meta: oligo-metachronous BM, and poly: poly metastasis), (negative: no stained cells, low: <10%, moderate:10-40%, and high: >40%)

### Slide 9
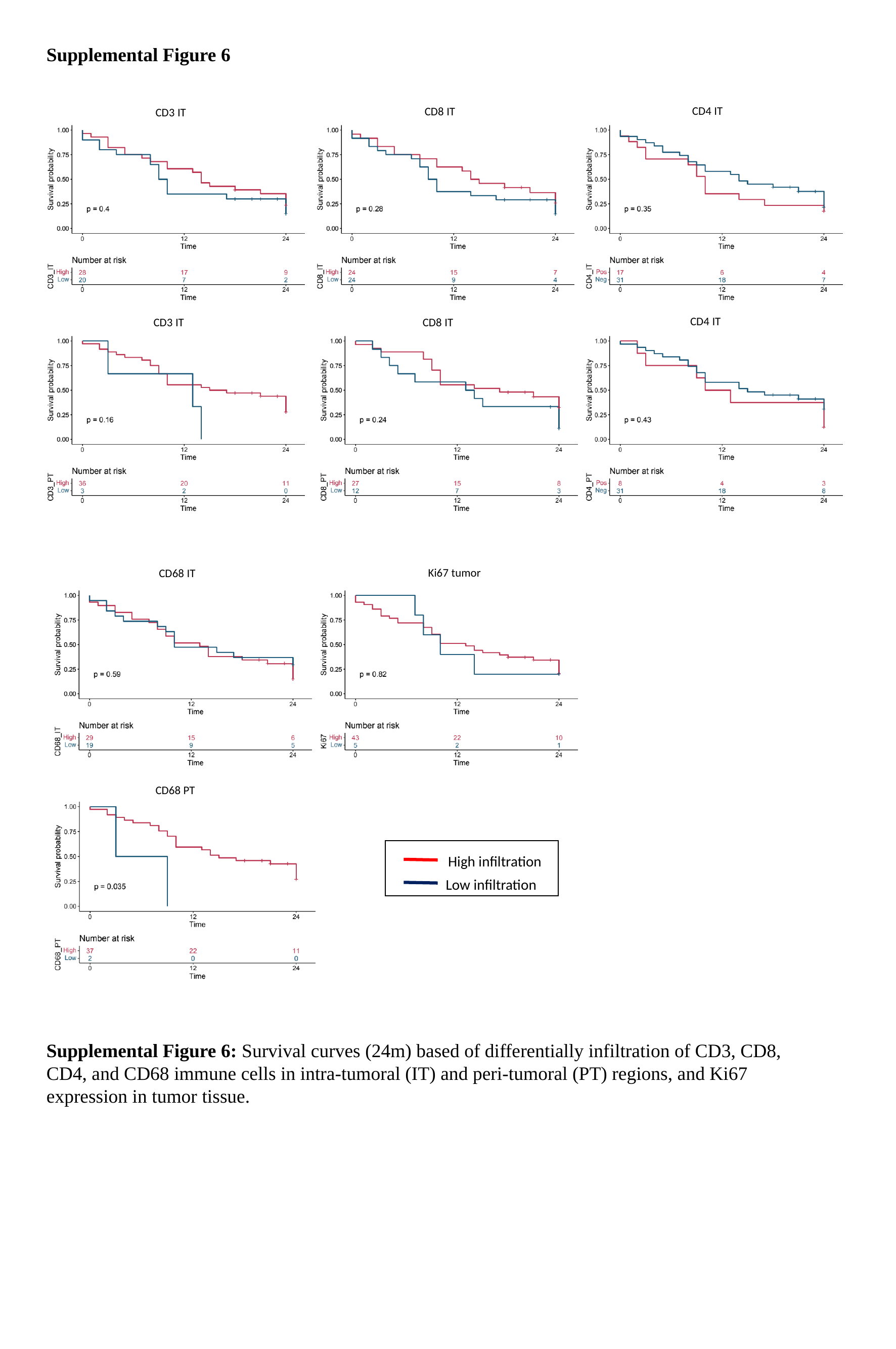

Supplemental Figure 6
CD4 IT
CD8 IT
CD3 IT
CD4 IT
CD8 IT
CD3 IT
Ki67 tumor
CD68 IT
CD68 PT
High infiltration
Low infiltration
Supplemental Figure 6: Survival curves (24m) based of differentially infiltration of CD3, CD8, CD4, and CD68 immune cells in intra-tumoral (IT) and peri-tumoral (PT) regions, and Ki67 expression in tumor tissue.

### Slide 10
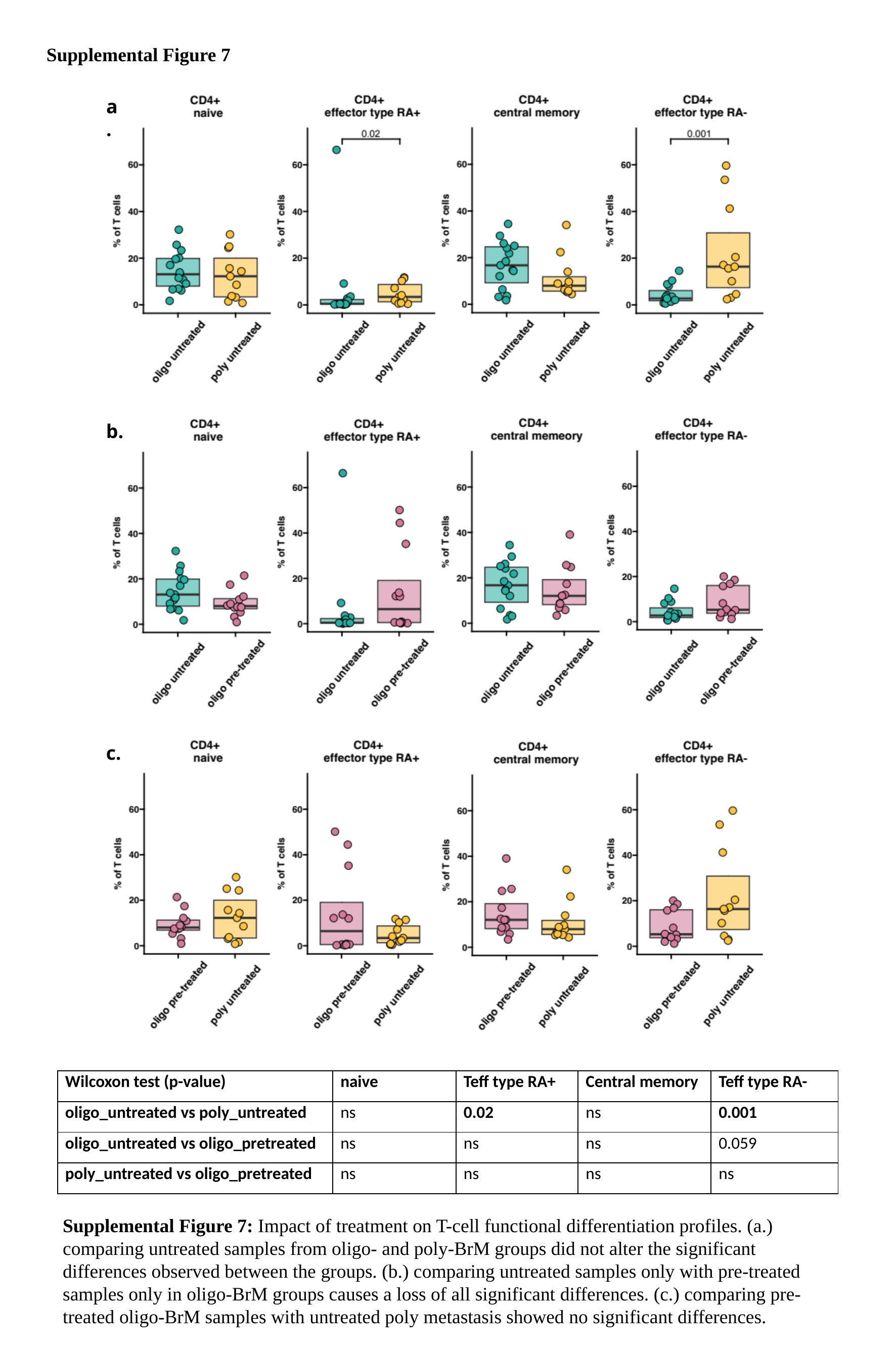

Supplemental Figure 7
a.
b.
c.
| Wilcoxon test (p-value) | naive | Teff type RA+ | Central memory | Teff type RA- |
| --- | --- | --- | --- | --- |
| oligo\_untreated vs poly\_untreated | ns | 0.02 | ns | 0.001 |
| oligo\_untreated vs oligo\_pretreated | ns | ns | ns | 0.059 |
| poly\_untreated vs oligo\_pretreated | ns | ns | ns | ns |
Supplemental Figure 7: Impact of treatment on T-cell functional differentiation profiles. (a.) comparing untreated samples from oligo- and poly-BrM groups did not alter the significant differences observed between the groups. (b.) comparing untreated samples only with pre-treated samples only in oligo-BrM groups causes a loss of all significant differences. (c.) comparing pre-treated oligo-BrM samples with untreated poly metastasis showed no significant differences.
